# Targeting Impaired Type I Interferon–IL-27 Signaling Rescues T Regulatory Cell Suppressive Function in Relapsing-Remitting Multiple Sclerosis

**DOI:** 10.1101/2025.08.26.671836

**Authors:** Manal Elzoheiry, Maryam Seyedsadr, John Wrobel, Yan Wang, Sowmya Shree Gopal, Ezgi Kasimoglu, Paulien Baeten, Ibrahim Hamad, Mia Rose Gleason, Bieke Broux, Jenny Ting, Booki Min, Silva Markovic-Plese

## Abstract

T Regulatory T cells (Tregs) from patients with relapsing-remitting multiple sclerosis (RRMS) exhibit impaired suppressive function, yet the underlying molecular mechanisms remain elusive. Single-cell RNA sequencing (scRNAseq) of *ex vivo-*sorted Tregs from RRMS patients and matched healthy controls (HCs) revealed down-regulation of type I IFN (IFN) and IL-27 signaling pathways in RRMS Tregs. These Tregs showed reduced expression of IFN-stimulated genes (ISGs) (*ISG15, MX1, IFITM1, IFI44L, OAS1*), as well as key mediators of Treg suppressive function (*LGALS3, CD81, FCRL3, CD7, CSTB*), all suggesting a key role of decreased IFN signaling in RRMS Treg dysfunction. To therapeutically target IFN signaling pathways and improve Treg suppressive functions, we used cGAMP-loaded microparticles (MPs) to activate the stimulator of IFN genes (STING) in experimental autoimmune encephalomyelitis (EAE). cGAMP-MP treatment ameliorated EAE via induction of Tregs expressing IL-27R, IL-10, TGF-b, and Granzyme B. This effect was abolished in Treg-specific IL-27R (Treg^ΔIl27ra^) knockout mice, confirming that IL-27 signaling is essential for Treg suppression.

*In vitro* IL-27 stimulation of RRMS-derived Tregs restored expression of IFN pathway genes (*IRF1, IFNGR, IFI16*) and Treg suppressive genes (*ICOS, IKZF3, IL7R, TIGIT*). Thus, we propose that IL-27 pre-stimulation may restore their suppressive function and migration (via *CCR6, CCR7, S100A11* and *S1PR4*) to the central nervous system (CNS) in future clinical trials.

**Significance:** Several studies have reported a role for type I IFN and IL-27 signaling in the induction of suppressive Tregs in autoimmune diseases. We report that RRMS Tregs have decreased expression of type I IFN and IL-27 signalling-related genes in comparison to HCs. The animal model of MS (EAE) was successfully treated with cGAMP-MPs, which, via induction of type I IFN, IL-27 and IL-10, restored Treg suppressive function. A scRNAseq study of Tregs from MS patients revealed that IL-27 *in vitro* stimulation normalized the expression of type I IFN genes and Treg suppressive genes. We propose that IL-27 pre-treatment may enhance Treg suppressive function and migration to the CNS in future clinical trials.

## Introduction

Patients with RRMS have deficient Treg cell suppressive function, which may contribute to disease pathogenesis (1). In this study, we used scRNAseq to determine underlying transcriptional changes causing decreased Treg suppressive function in RRMS. scRNAseq of sorted CD4^+^CD25^+^CD127^low^ *FOXP3^+^* activated Tregs from untreated RRMS patients and matched HCs (activated Tregs clustered based on *FOXP3, IL2RA, ID2, CCR2, ID2, S100A4, S100A6, ICOS, ITGB1, CXCR3, DUSP2, NR4A1, EGR1, and IKZF2* expression) (2) revealed that response to type I IFN and IL-27 signaling pathway are among the most down-regulated pathways in MS Tregs. The results suggest that impaired endogenous type I IFN/IL-27 signaling is a key regulator of Treg suppressive function. The differentially expressed gene (DEG) analysis revealed significantly decreased type I IFN (*ISG15, MXI, IFITM1, IFI44L, OAS1*) and Treg suppressive function (*LGALS3, CD81, FCRL3, CD7, CSTB)* genes in RRMS Tregs. The results support a causative role of decreased IFN signaling in dysfunctional Treg suppression in MS. Consistent with our prior report (3), RRMS patients had significantly decreased CSF and serum IFN-β and IL-27 levels in comparison to HCs.

To therapeutically target deficient type I IFN function in Tregs in the animal model of the MS, EAE, we used MP-delivered cGAMP, a Stimulator of Interferon Genes (STING) agonist, which inhibited EAE via induction of type I IFN, IL-27, and IL-10-secreting tolerogenic DCs. The therapeutic effect was completely reversed in *Il27ra ^-/^*^-^ mice (4), suggestive of IL-27 induction as a key mediator of the therapeutic effect. Since IL-27 was reported to inhibit EAE only in the presence of Tregs (5), we examined whether cGAMP-MP-induced IL-27-secreting DCs suppress the disease via induction of Tregs. The treatment-induced CNS tolerogenic CD80/CD86^low^IL-27^+^ DCs increased the frequency and suppressive function of the CNS-infiltrating FoxP3^+^ Tregs. These suppressive Tregs expressed IL-27R, IL-10, TGF-β, and Granzyme B. The loss of cGAMP-MP treatment effect in Treg-specific *IL-27R*^-/-^ (Treg^ΔIL27ra^) mice (6) revealed that IL-27R signaling is required for Treg suppressive function.

Following reports that IL-27-stimulated Tregs have enhanced suppressive function in mouse model of colitis (7), we examined whether *in vitro* IL-27 stimulation can restore impaired Treg suppression in RRMS. scRNAseq analysis of sorted Tregs following *in vitro* IL-27 stimulation revealed an increased (reconstituted) type I and II IFN/IL-27 signaling-related genes *IRF1, IF16* and *IFNGR1*. Since IL-27 treatment also restored the expression of genes that mediate Treg suppressive function (*ICOS, IKZF3, IL7R*, and *TIGIT*), we propose that IL-27 pre-stimulation of autologous Tregs can be used prior to their therapeutic infusion. The autologous Treg transfer is currently tested in clinical trials aiming to reconstitute immune tolerance in RRMS (8) and other autoimmune diseases (9).

## Results

### RRMS Tregs have decreased expression of type I IFN and IL-27 signaling pathways and of Treg suppressive genes

To characterize transcriptome changes in Tregs from RRMS patients, we performed scRNAseq of sorted CD4^+^CD25^+^CD127^low^ Treg cells from three untreated RRMS patients (10) and three age-, sex-, and race-matched HCs (Fig 1A, SI Appendix, Table S1).

**Figure 1.**
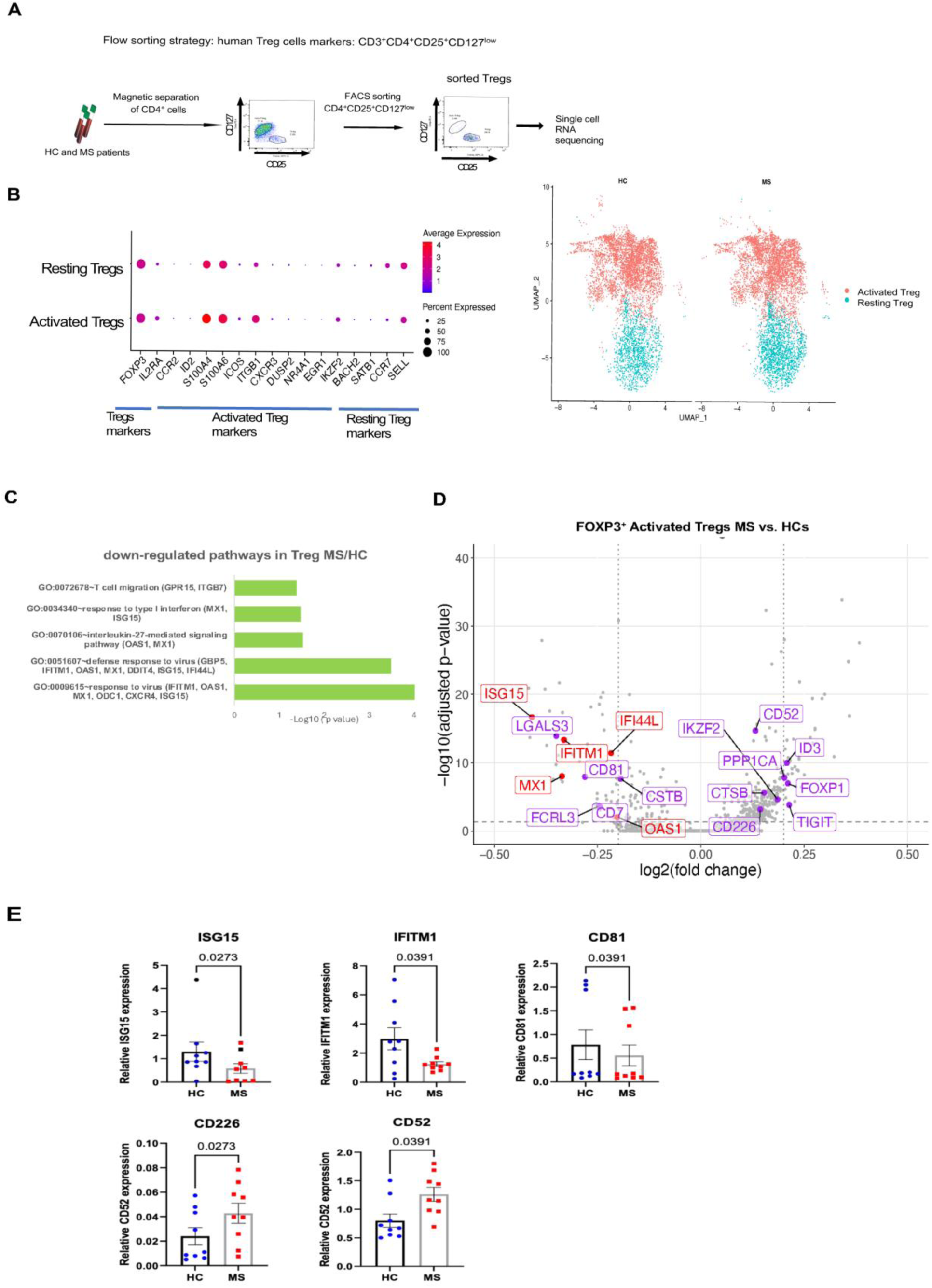
scRNAseq analysis of Tregs from RRMS patients revealed decreased expression of type I IFN and Treg suppressive genes. (*A*) Schematic representation of the experiment. Magnetic bead-separated CD4^+^ cells from three untreated RRMS patients and three matched HCs were sorted for CD4^+^CD25^+^CD127^low^ Treg cells and submitted for scRNAseq experiment. (*B*) Dot plot showing cluster-defining gene expression in activated and resting *FOXP3^+^* Tregs. UMAP clustering of activated and resting *FOXP3^+^* Tregs was derived from the initial 8 clusters (*SI Appendix,* Fig. 1 *B*). (*C*) Gene Ontology and pathway enrichment analysis of DEGs in activated *FOXP3^+^* RRMS Tregs versus HC subjects. (*D*) Volcano plot showing biologically relevant DEGs in RRMS Tregs compared to HCs. Color codes: type I IFN (red), and Treg suppressive genes (purple). DEGs with adjusted Log10p<0.05 and Log2FC>0.1. *(E)* RT-PCR validation of selected DEGs identified by scRNAseq experiment was completed in sorted Treg from 10 untreated RRMS patients and matched HCs. Data are presented as relative gene expression; statistical analysis was performed using Wilcoxon test.

After quality control check and filtering, a total of 61,390 cells were analyzed: 28,971 from HCs and 32,419 from RRMS patients *(SI Appendix, Fig.* S1*A).* We analyzed *FOXP3^+^* Tregs (11,643 cells), which constituted 19% of total sorted cells, consistent with previous reports (11). Eight clusters of *FOXP3^+^* cells *(SI Appendix, Fig.* S1*B)* were grouped into *activated* and resting Treg cells based on the gene expression for activated Tregs (*FOXP3, IL2RA, CCR2, ID2, S100A4, S100A6, ICOS, ITGB1, CXCR3, DUSP2, NR4A1, EGR1, IKZZF2*) and resting Tregs (*BACH2, SATB1, CCR7, SELL*) (2). Since the initial analysis did not show difference between MS and HC Treg clusters, and DEG analysis revealed 395 DEGs in activated and only 8 in resting Tregs, we focused our analysis on activated or memory Tregs, which are reported to have suppressive function (2).

The absolute numbers (and percentages) of activated (4,266 and 4,153); and resting Tregs (1,870 and 1,340) did not differ between MS and HCs, and the two Treg subsets represented distinct populations (Fig. 1*B*). Among the 395 DEGs in *activated Tregs,* 93 were down-regulated and 301 up-regulated in RRMS Tregs compared to matched HCs (Dataset S1).

Pathway enrichment analysis of down-regulated DEGs in activated Tregs (2) revealed strong enrichment for type I IFN and IL-27 signaling pathways (Fig 1*C*, Dataset S2). To characterize the molecular mechanisms of type I IFN regulation of Treg suppressive function, we manually curated biologically relevant DEGs and grouped them into six functional groups: (1) genes involved in type I IFN signaling; (2) cell death/apoptosis; (3) cell migration to the inflammatory sites; (4) cytokine signaling; (5) T cell receptor (TCR) signaling; and (6) Treg suppressive function (Dataset S1). Fig. 1*D* shows a volcano plot highlighting annotated type I IFN signaling genes (red) and Treg suppressive genes (purple), while a volcano plot in *SI Appendix, Fig.* S1*C* shows all biologically relevant Treg DEGs with -Log10 adjusted p value <0.05 and Log2 fold change (FC) >0.1.

<u>Down-regulated DEGs in RRMS activated Tregs</u> included *ISG15, MXI, IFITM1, IFI44L, and OAS1,* which are involved in type I IFN signaling (12), as well as Treg suppressive function genes (*LGALS3, CD81, FCRL2, CD7, and CSTB*). These two groups are of particular interest, as type I IFN signaling is proposed to regulate Treg suppressive function (12–14). ISG15 is typically increased in activated and IFN-stimulated Tregs and can induce IFN-γ and IL-10 secretion (15), which may, in a positive feedback fashion, promote suppressive Treg activity. However, ISG15 also negatively regulates responsiveness to type I IFN by blocking STAT1 signaling (16). Following our report on decreased type I IFN signaling in CD4^+^ cells from MS patients (3), we propose that decreased *ISG15* expression reflects attenuated type I IFN signaling in RRMS Tregs. Down-regulated DEGs associated with Treg suppressive function include: *LGALS3,* which encodes Galectin-3, a Treg cell marker (17); *CD81,* a costimulatory molecule (18); *CD7*, required for Treg homeostasis; and *FCRL3,* required for optimal Treg function (19). Selected biologically relevant gene expression levels are presented in violin plots *(SI Appendix,* Fig. S2).

Up-regulated DEGs in RRMS activated Tregs included Treg suppressive genes: *TIGIT,* a co-inhibitory molecule that selectively inhibits Th1 and Th17 responses (20); *CD226,* which competes with TIGIT for their common ligand CD155, and negatively affects Treg suppression and proliferation (21, 22)*; Foxp1*, a transcription factor critical for maintaining Treg suppressive function (23); *ID3,* maintaining the Treg pool (24); and the costimulatory Treg gene *CD52* (25, 26).

To validate the scRNAseq data, we analyzed sorted CD3^+^CD4^+^CD25^+^CD127^low^ Tregs from an independent cohort of 10 untreated RRMS patients and 10 age-, sex-, and race-matched HCs (*SI Appendix, Table S1*). We confirmed decreased gene expression in RRMS Tregs for type I IFN-induced genes *ISG15* (−2.1-fold change, p=0.027) and *IFITM1* (−2.3-fold, p=0.039), and for the costimulatory/migration molecule *CD81* (−1.4, p=0.039). Validation of up-regulated genes revealed increased expression of *CD226* (1.8-fold, 0.027) and *CD52* (1.6-fold, p=0.039) genes (Fig. 1 *E*). Together, these data suggest that impaired type I IFN signaling may contribute to deficient Treg suppression in RRMS.

### IFN-β and IL-27 levels are decreased in the CSF and serum of RRMS patients

Following our findings of decreased type I IFN and IL-27-induced gene expression in RRMS Tregs, we measured IFN-β and IL-27 levels in CSF and serum samples from untreated RRMS patients and HCs (CSF 15 MS, 20 HC; serum 20 MS, 9 HC) (*SI Appendix, Table S2*). We found significantly decreased levels of IFN-β and IL-27 in the CSF of RRMS patients, as well as reduced IFN-β serum level compared to controls (Fig. 2 *A*). A positive correlation between serum IFN-β and IL-27 levels (Fig. 2 *B*), likely reflects IFN-β-induced IL-27 secretion (27).

**Figure 2.**
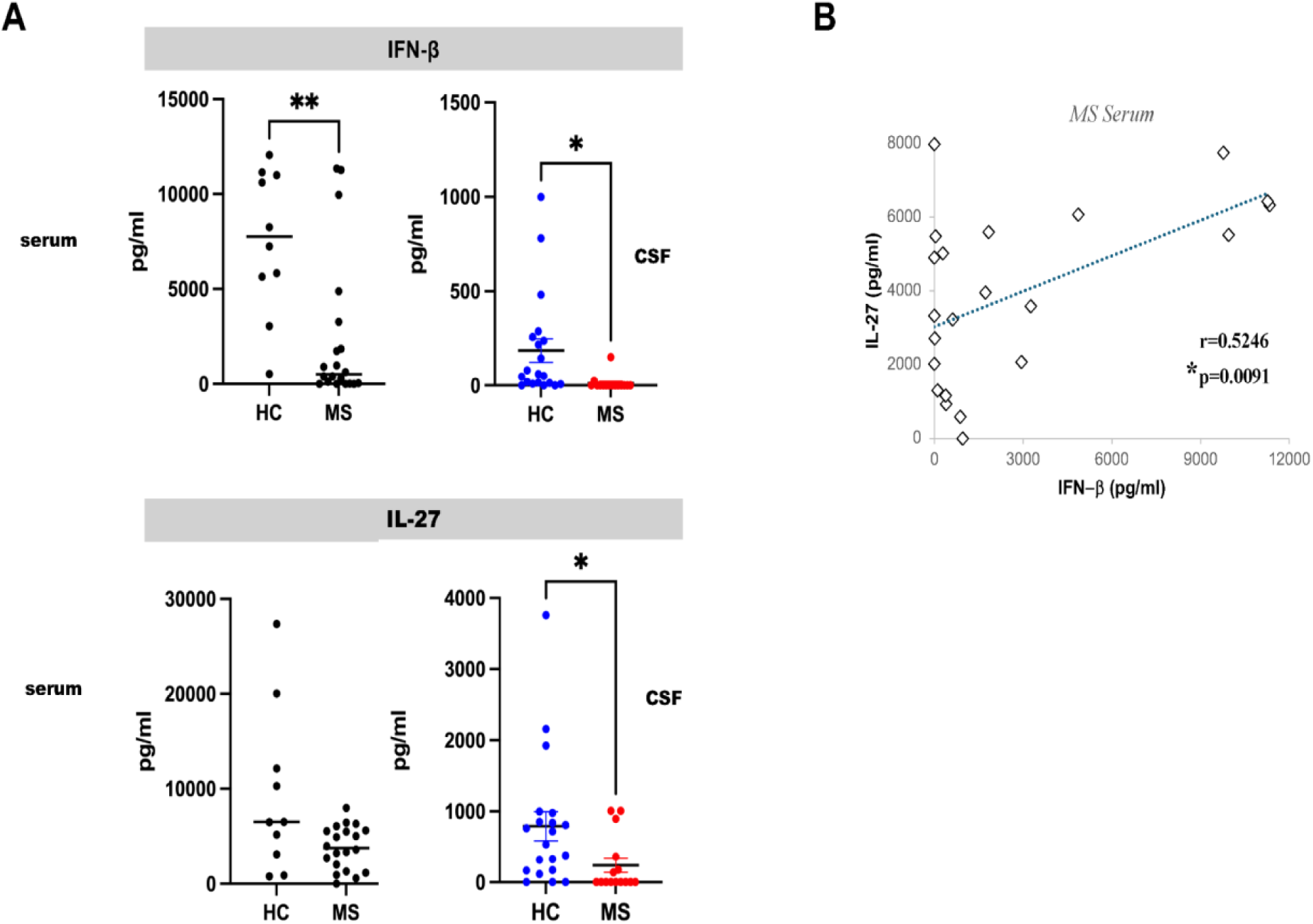
IFN-β and IL-27 levels are decreased in the CSF and serum from RRMS patients. *(A)* IFN-β and IL-27 levels in CSF (15 MS and 20 HC samples) and serum (20 MS and 9 HC samples) were compared using Mann-Whitney test. (*B*) Positive correlation between IFN-β and IL-27 serum levels in RRMS patients, Spearman correlation.

### cGAMP-MPs suppress RREAE by IL-27-induced suppressive Tregs

To determine the extent to which stimulation of endogenous type I IFN and IL-27 secretion by myeloid cells may normalize Treg suppressive function in EAE, we treated mice with cyclic G(3’,5’)pA(3’,5’)p (cGAMP), STING agonist encapsulated in MPs. cGAMP is delivered intracellularly via MPs, which are phagocytosed by myeloid cells (monocytes and DCs).

Our previous study reported that cGAMP-MPs suppressed EAE by the induction of tolerogenic DCs that secreted IFN-β, IL-27 and IL-10 (4). In the present study, we tested whether the therapeutic effect of cGAMP-MPs is mediated by the induction of suppressive Tregs.

We immunized 8–10-week-old female *SJL/J* mice with proteolipid protein (PLP)139-151 peptide to induce RREAE. Mice were divided into two groups: the first group received intramuscular (I.M.) injections of cGAMP-MPs (5 μg/mouse), and the second group received blank-MPs. The treatment started either at the onset of clinical symptoms (day 9 post-immunization (p.i.) or at the peak of disease (day 15 p.i.), with a total of 5 injections. Fig. 3 *A* and *SI Appendix, Fig. S*4A show that cGAMP-MP treatment significantly decreased clinical scores in mice treated at both time points. Plasma levels of IL-27 and IL-10 were increased in both treatment groups (Fig. 3 *A* and *SI Appendix, Fig. S*4A).

**Figure 3.**
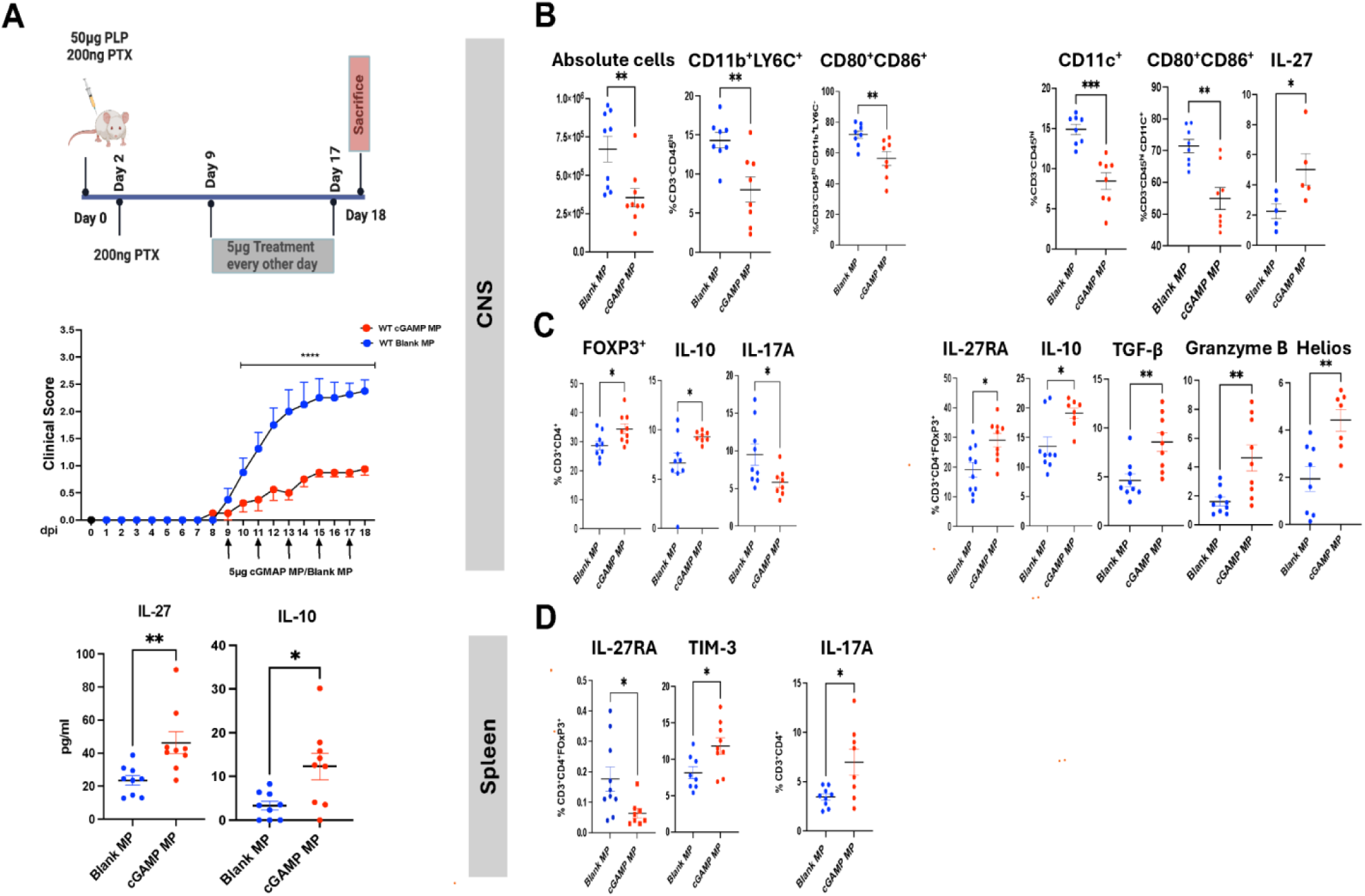
cGAMP-MPs reduce RREAE disease severity via IL-27-induced suppressive Tregs. (*A)* SJL/J mice with RREAE were treated with cGAMP- or blank-MP (5 μg/mouse) starting on day 9 (disease onset, 13 mice/group, *left panel*) and then every other day for five doses. Mice were scarificed on day 24 p.i. Clinical scores were monitored daily. Statistical analysis was performed using two-way ANOVA. Data are presented as mean ± SEM. **** denotes p<0.0001. Serum cytokines were measured in 9 mice/group by ELISA. Data are presented as mean ± SEM; statistical analysis was performed using Mann–Whitney test. *p<0.05, ** p<0.01. (*B)* Flow cytometry analysis of CNS-infiltrating inflammatory cells (9 mice/group): the percentages of Ly6C^+^ monocytes and CD11c^+^ DCs and their expression of CD80, CD86, and IL-27. *(C)* The frequency of FoxP3^+^, IL-10^+^, and IL-17A^+^ CD3^+^CD4^+^ cells within CNS infiltrates were determined. IL-27RA^+^, IL-10^+^, TGF-β^+^, granzyme-B^+^ and Helios^+^ cells were determined within CNS CD3^+^CD4^+^FoxP3^+^ Tregs. *(D)* In the spleen, the frequencies of CD3^+^CD4^+^FoxP3^+^ cells expressing IL-27RA and TIM-3, and CD3^+^CD4^+^IL-17A^+^ cells, were compared between blank-MP and cGAMP-MP treated groups. Data are presented as mean ± SEM; the Mann–Whitney test was used for comparison. *p<0.05, ** p<0.01, *** p<0.001.

Flow cytometry analysis of CNS infiltrates from mice treated at the onset of disease revealed reduced numbers of inflammatory cells and decreased percentages of CNS-infiltrating CD11b^+^Ly6C^+^ monocytes and CD11c^+^ DCs (gating strategy presented in *SI Appendix, Fig. S3*). Treatment also decreased their expression of costimulatory CD80 and CD86 and increased their IL-27 production (Fig. 3 *B*), consistent with tolerogenic DC phenotype (4). Importantly, we detected an increased frequency of FoxP3^+^CD4^+^ Tregs in CNS infiltrates, with elevated expression of IL-27R, IL-10, TGF-β, and Granzyme B, and a reduced frequency of pathogenic IL-17A^+^CD4^+^ cells in cGAMP-MP-treated mice (Fig. 3 *C*). Similarly, there was an increased percentage of CD3^+^CD4^+^FoxP3^+^ IL-27R^+^ and TIM-3^+^ Tregs in the spleens of cGAMP-MP-treated mice (Fig. 3 *D*), as well as in the lymph nodes (LNs) from mice treated at the peak of the disease (*SI Appendix, Fig. S*4C). These results suggest that type I IFN and IL-27 signaling promote the *in vivo* expansion of CNS FoxP3^+^IL-27R^+^ Tregs with enhanced suppressive function.

To mechanistically support these *in vivo* findings, we treated magnetic bead-separated CD11b^+^Ly6C^+^ monocytes *in vitro* with cGAMP-MPs, which led to decreased CD80 and CD86 surface expression and increased secretion of IL-27 and IL10 (*SI Appendix, Fig. S*5 *A, B*). Co-culture of pretreated monocytes with CD4^+^ T cells for 48 h further elevated IL-27 and IL-10 levels in supernatants and increased the percentage of CD4^+^FoxP3^+^ Tregs expressing GITIR within the co-cultured CD4^+^ cells (*SI Appendix, Fig. S*5 C).

Finally, we demonstrated increased expression of type I IFN-regulated genes *ISG15, OAS1* and *IFITM1* in magnetic-bead purified spleen Tregs from cGAMP-MP-treated mice at both the onset and peak of the disease (*SI Appendix, Fig. S*6). These results validate the above scRNAseq results from RRMS patients showing decreased expression of those genes in RRMS Tregs (Fig. 1 *E*).

### IL-27R signaling in Tregs increases their suppressive function in EAE

As previously reported by our group (6), mice with Treg-specific *IL27R* deletion (Treg^ΔIl27ra^) exhibited loss of Treg function and worsened clinical scores in EAE (*SI Appendix, Fig. S*7A). Flow cytometry analysis of CNS infiltrates of Treg^ΔIl27ra^ mice with EAE revealed decreased percentages of CD4^+^FoxP3^+^ Tregs expressing IL-27R, TGF-β, and Granzyme B, and increased percentages of IL-17A^+^ and TNF^+^ effector CD4^+^ T cells, compared to untreated wild-type (WT) mice with EAE (*SI Appendix, Fig. S*7B-C). In the spleen of the same untreated Treg^ΔIl27ra^ mice, we observed decreased percentages of LAG3^+^, TIGIT^+^, GITR^+^, IL-10^+^ and TGF-β^+^ Tregs (*SI Appendix, Fig. S*7D), indicating that *in vivo* IL-27R signaling induces those co-inhibitory molecules critical for Treg suppression (6, 7).

To directly test the role of IL-27R signaling in regulating Treg suppressive function, we treated Treg^ΔIl27ra^ mice in comparison to WT mice with cGAMP-MP or blank-MP at the onset of EAE (Fig. 4 *A*). The therapeutic effect of cGAMP-MPs was abolished completely in Treg^ΔIl27ra^ mice. They had significantly decreased frequencies of the CNS CD4^+^FoxP3^+^ Tregs expressing CTLA-4, GITR, PD-1, and TGF-β co-inhibitory markers in comparison to treated WT mice (Fig. 4 *B*), suggesting impaired migratory capacity of IL27R^-^/^-^ Tregs across the blood-brain barrier (BBB). In the spleens of cGAMP-MP-treated Treg^ΔIl27ra^ mice, we found decreased percentages of LAG3^+^ and CTLA-4^+^ Tregs. These findings suggest that IL-27-LAG3 axis induces CTLA-4 expression, which is required for optimal Treg suppressive function (6, 28). Additionally, the proliferation of Tregs, measured by intracellular Ki67 expression, was decreased in treated Treg^ΔIl27ra^ mice, indicating that IL-27 is required for Treg expansion (Fig. 4 *C*). These results demonstrate the critical role of IL-27R signaling in Treg suppressive function.

**Figure 4.**
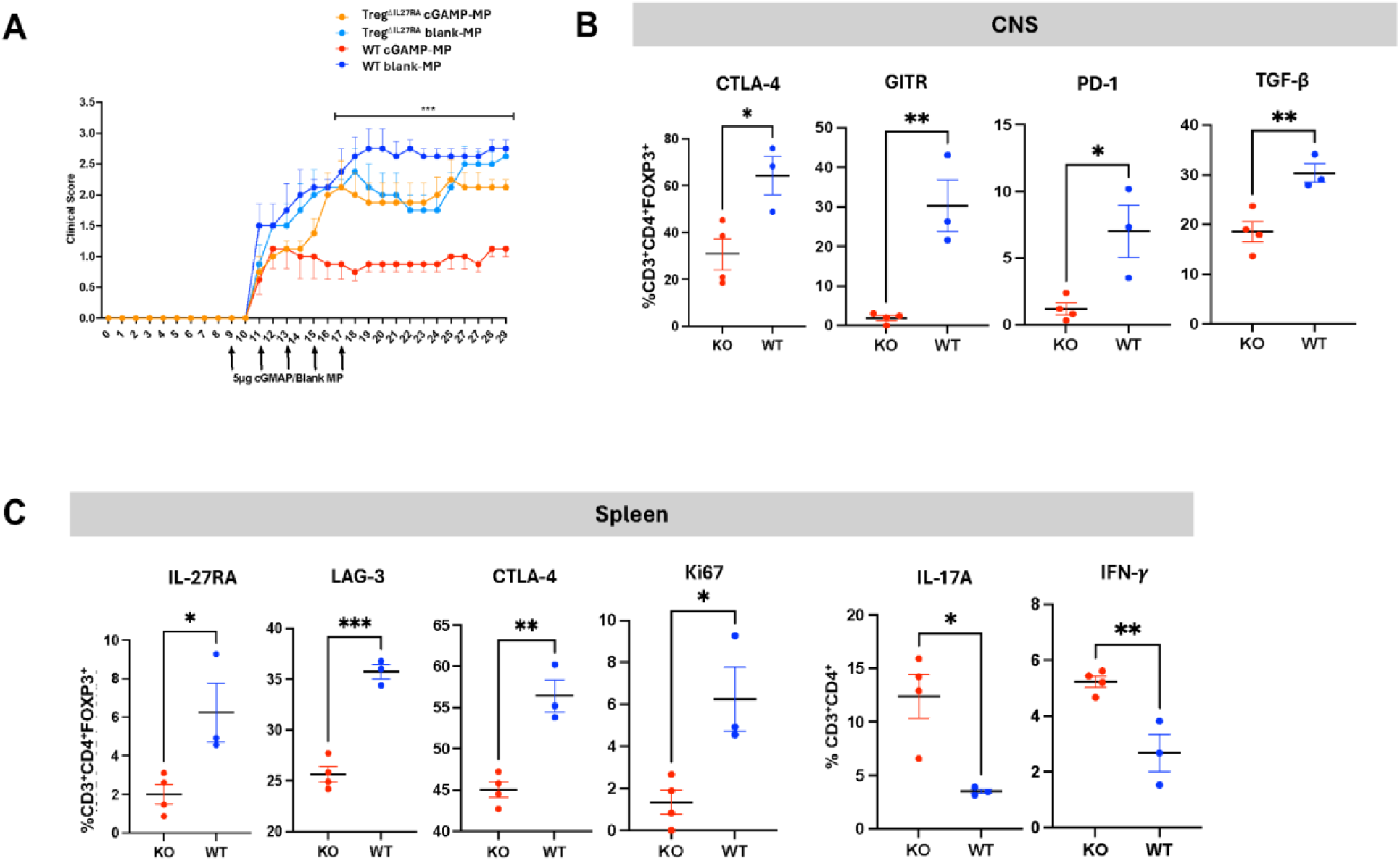
cGAMP therapeutic effect is mediated via IL-27-induced Treg suppressive function. *(A)* The therapeutic effect of cGAMP-MPs is abolished in Treg^ΔIL27ra^ mice. Data are presented as mean±SEM; statistical analysis was performed using two-way ANOVA, *** p<0.01. *(B)* Treg^ΔIL27ra^ and WT mice (3-4/group) treated with cGAMP-or blank-MP beginning on day 9 were sacrificed on day 18. The frequency of CNS-infiltrating FoxP3^+^ Tregs expressing CTLA-4, GITR, PD-1, and TGF-β were assessed. *(C)* In the spleens of cGAMP-MP-treated Treg ^ΔIL27ra^ and WT mice, the expression of IL-27RA, LAG3, CTLA-4, and proliferation marker Ki67 was measured in CD3^+^CD4^+^FOXP3^+^ cells. Data were presented as mean±SEM and statistical analysis was performed using the Mann–Whitney test. *p<0.05, and **p<0.01.

### IL-27 normalizes the expression of IFN and Treg suppressive genes in RRMS Tregs

A second scRNAseq experiment on sorted CD4^+^CD25^+^CD127^low^ Tregs from four untreated RRMS patients and age-, sex-, and race-matched HC donors (*SI Appendix, Table S3*) was performed to test transcriptome changes in MS and HC Tregsu after IL-27 *in vitro* stimulation. In this analysis we focused on *FOXP3^+^* Tregs. Fig. 5 *A* presents pathway analysis of 473 DEGs between CD4^+^CD25^+^CD127^low^ sorted *FOXP3* gene-expressing Tregs from RRMS and HC donors, highlighting the down-regulation of type I IFN production as one of the most suppressed pathways (Dataset 3 and 6). Fig. 5 *B* presents DEGs in a volcano plot: 390 down-regulated (adjusted p value <0.05, and Log2FC<0.25) and 83 up-regulated genes in RRMS Tregs. We manually curated the list of DEGs detected in sorted *FOXP3^+^* Tregs as in Fig 1. Volcano plot presents significantly <u>down-regulated DEGs,</u> including IFN signaling (*IRF1, IFI16,* and *IFGNR1*), (red) and Treg genes that regulate suppressive function (*ICOS, IZKF3, IL7R, TIGIT, and TRIM13*) (purple). Decreased DEGs in RRMS Treg also contained migratory genes (*CCR4, CCR6, CCR7,* CXCR6*, ICAM2, ITM2C, S100A11, S1PR1, S1PR4*) (yellow); cytokine secretion genes (*MAP3K2, NFKB1, NFKB2, REL, RELB, TNFRSF18, TNFRSF9, TRAF1, TRAF3*) (green); and multiple TCR signaling *(CD247)* genes (blue) (Supplemental Dataset S3). <u>Up-regulated genes</u> include: IFN-induced antiviral gene *IFITM2* (29) with pro-apoptotic effect (30); and *PRDM1*, recently proposed key mediator of dysfunctional Tregs in RRMS via inhibition of IFN signaling (31). *PRDM1* encodes BLIMP-1 protein, a transcriptional repressor that binds the IFNB1 promoter and suppresses its expression (32). Treg suppressive genes include costimulatory *CD52* (33), *CD69* marker of tissue resident Tregs (34), and *IL32* which promotes suppressive Treg function (35). Additional up-regulated genes contain cell death-related *BCL2* and *FOS*; *LAIR2* regulator of T cell activation (36); and a suppressor of cytokine secretion *SOCS3*.

**Figure 5.**
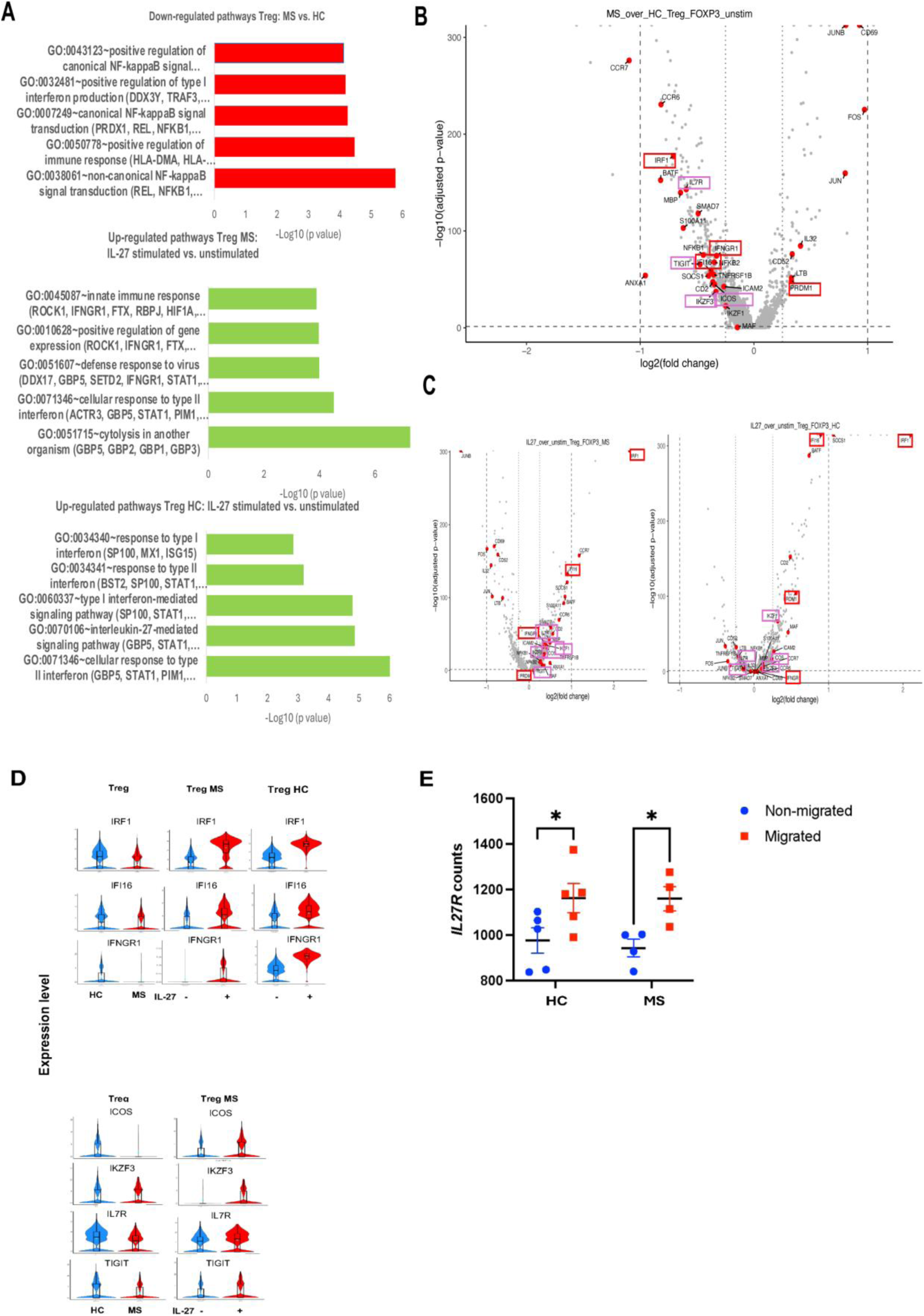
IL-27 normalizes the expression of IFN and Treg suppressive genes in RRMS. (*A*) Gene Ontology and pathway enrichment analysis of DEGs from *FOXP3^+^* Tregs from RRMS versus HC subjects; MS Tregs IL-27-stimulated vs. unstimulated; and HC Tregs IL-27-stimulated vs. unstimulated. (*B*) Volcano plot showing biologically relevant DEGs in RRMS Tregs compared to HCs. Color codes: type I IFN (red), and Treg suppressive genes (purple). *(C)* Volcano plot showing biologically relevant DEGs in RRMS Tregs IL-27-stimulated vs. unstimulated cells (left) and HC Tregs IL-27-stimulated vs. unstimulated (right), DEGs with adjusted Log10p<0.05 and Log2FC>0.25 and Log2FC<0.25. *(D)* Violin plots present gene expression level of IFN genes decreased in MS Tregs vs HC Tregs, which are upregulated after IL-27 stimulation of MS Tregs and HC Tregs (upper panel). Volcano plots present selected Treg suppressive gens that are decreased in MS Tregs and up-regulated following IL-27 stimulation. *(E) IL27R* gene expression in migrated vs. non-migrated Tregs from RRMS and HC donors following 24 h migration assay. Statistical analysis using two-way ANOVA.

We next tested the transcriptome changes of the same sorted Tregs stimulated *in vitro* with IL-27 (1 hour (h), 25 ng/ml). scRNAseq of <u>IL-27-stimulated vs. unstimulated</u> <u>RRMS</u> Tregs revealed <u>up-regulation</u> of type I IFN (*IFI6, IFIT2, IFIT3, IFNGR1, IRF1, IRF4, IRF9*, and *STAT1*) and Treg suppressive genes (*ICOS, IZKF1, IZKF3, IZKF4, IL2RA, IL2RB, IL7R,* and *TIGIT*), (Fig. 5 *B-D*, Dataset 4 and 7). IL-27 also increased the expression of migration-related genes (*CCR6, CCR7, ICAM2, S100A11* and *S1PR4*).

Notably, multiple genes with decreased expression in RRMS in comparison to HC Tregs were normalized in MS Tregs upon IL-27 stimulation, including IFN-related genes (*IRF1, IFI16,* and *IFNGR1*), and Treg suppressive genes (*ICOS, IKZF3, IL7R* and *TIGIT*); migratory genes (*CCR6, CCR7, S100A11* and *S1PR4*) and inhibitory cytokine related genes (*SMAD7* and *SOCS1*). Those changes support the requirement of IL-27R signaling for Treg suppressive function, migration, and expansion, potentially mediated via activation of type I and II IFN signaling (Fig. 5 *D* and Dataset 4, last sheet provides a list of overlapping genes between MS Treg down-regulated and IL-27 stimulation-restored genes in MS Tregs).

The final observation following comparison of <u>IL-27-stimulated Tregs from RRMS</u> <u>and HCs</u> donors is that multiple type I IFN-up-regulated genes (*IFI6, IFIT2, IFIT3, IRF1, IRF4*, and *STAT1*) in RRMS Tregs did not reach the level of expression as in HC Tregs, indicating impaired IL-27-induced IFN signaling in RRMS. This was also noted for the Treg genes (*IKZF1, IKZF4* and *IL2RA*), reinforcing the idea that deficient IL-27/IFN signaling in MS can be partially restored upon IL-27 stimulation (Fig. 5 *C (right)*, and Dataset 5). Fig. 5 *D* presents the expression levels of IFN genes (*IRF1, IFII6, IFNGR1*) in MS vs. HC Tregs in violin plots, whose decreased expression levels in MS Tregs is normalized upon IL-27 stimulation. The lower panel presents the key Treg suppressive genes (*ICOS, IZKF3, IL7R* and *TIGIT*), which were decreased in MS Tregs compared to HC, and reconstituted upon IL-27 stimulation. Datasets S3 shows a list of overlapping genes that are decreased in MS Tregs and increased upon IL-27 stimulation. Dataset 7 and 8 present pathway analysis of the IL-27-stimulated vs. non-stimulated RRMS Tregs, and HC Tregs, respectively.

To address the role of IL-27R signaling in Treg capacity to migrate to the CNS, we analyzed the *in vitro* migrated vs. non-migrated sorted Tregs from RRMS and HC donors across a human endothelial cell barrier (17). The RNAseq study of Tregs showed an increased *IL27R* gene expression in migrated vs. non-migrated Tregs from MS patients, while Tregs from HCs did not show a difference in *IL27R* gene expression (Fig. 5 *F*).

## Discussion

scRNAseq data analysis identified decreased expression of multiple type I IFN and IL-27 signaling genes in sorted CD4^+^CD25^+^CD127^low^ Tregs from RRMS patients, which may contribute to impaired Treg suppressive function in RRMS and other autoimmune diseases (37). We propose that reduced type I IFN and IL-27 signaling may regulate deficient Treg suppressive functions in RRMS and may represent a therapeutic target to optimize Treg suppressive function prior to using autologous Tregs therapeutically (31, 38).

The literature on type I IFN regulation of Treg function remains controversial. In chronic viral infections, type I IFN signaling can impair Tregs suppressive function (39, 40). In contrast, during chronic inflammation, Tregs lacking IFNAR exhibit compromised suppressive functions, leading to worsened inflammation (41–44). These findings suggest that the effects of type I IFN on Treg function are context dependent. Recent studies in autoimmune diseases report that type I IFN signaling enhances Treg suppressive function in RRMS by orchestrated regulation of co-inhibitory markers in Treg and Tconv (31). In a model of graft-versus-host disease (GVHD) disease, IFN treatment induces FoxP3 acetylation and stabilizes its expression, promoting sustained Treg function (12).

Studies investigating the role of IL-27 in regulation of Treg suppressive function have shown that systemic IL-27 administration prevents and attenuates ongoing EAE (45) and that IL27R ^-^/^-^ mice develop more severe EAE (46). Since Tregs numbers in the CNS of these mice were comparable to controls, it was concluded that they had a deficient suppressive function (5). Treg^ΔIL27Ra^ mice with specific Treg deletion of IL27R had severe EAE, similar to the systemic IL27R deletion. These findings indicate that IL-27R signaling in FoxP3^+^ Tregs is indispensable for their suppressive function and the control of EAE. Lag3 is induced in Tregs following IL-27 stimulation and inhibits inflammatory responses by suppressing DC maturation by contact-dependent engagement of its ligand MHC class II.

Immune cell functions are tightly linked to their metabolic programs (47, 48). Tregs rely on energy-efficient metabolic programs such as oxidative phosphorylation (OXPHOS) and fatty acid oxidation (FAO) to maintain their suppressive capacity (49). Glycolysis regulates Treg expansion and migration (50, 51); however, excessive glycolytic activity can impair their suppressive abilities (52, 53). Regarding IL-27’s role in regulating Treg metabolic program, our group recently demonstrated that the inhibitory co-receptor Lag3 enhances FoxP3^+^ Tregs suppressive function by inhibiting Myc-dependent glycolytic metabolic programming (28).

Several studies have emphasized the critical roles of type I IFN (31) and IL-27 for the induction of Treg suppressive function (5, 6), through induction of coinhibitory receptors CTLA-4, PD-1, TIM-3, LAG3 and TIGIT (54–56). Consistent with this, we observed decreased expression of LAG-3, TIGIT and GITR in Treg^ΔIl27ra^ mice with EAE (31, 56).

cGAMP MPs suppress RREAE by inducing tolerogenic monocyte-derived DCs that secrete IL-27 and IL-10, promoting expansion of IL-27R^+^Treg in the CNS (Fig. 3 *A*). *In vitro* co-culture of cGAMP MP-pretreated monocytes with CD4^+^ T cells confirmed expansion of FOXP3^+^ GITIR^+^ Tregs (*SI Appendix, Fig. S*5 C). In contrast, Treg^ΔIl27ra^ mice with Treg-specific *IL27ra* deletion did not respond to cGAMP-MP treatment, indicating that IL-27 signaling is required for the expression of LAG-3, CTLA-4, and GITR inhibitory markers and for the Treg suppressive function (Fig. 4 B, C).

Given that tolerogenic DC have decreased expression of costimulatory molecules and increased type I IFN, IL-27 and IL-10 secretion, they may support Treg expansion and are considered a promising treatment approach for RRMS (57). However, autologous Treg infusion remains a more direct method to reconstitute immune tolerance and is currently being tested in clinical trials (8).

Several Phase I clinical trials in autoimmune diseases (e.g., type 1 diabetes) and in transplant recipients have demonstrated the feasibility of expanding autologous Tregs and documented good tolerability of the intravenously administered autologous Tregs (9). Current efforts aim to improve Treg expansion, suppressive function, engineered TCRs for tissue homing, or use of chimeric antigen receptor (CAR) Tregs, and survival by using low-dose IL-2, (58).

Based on the presented data and previously demonstrated IL-27 pretreatment-induced improvement of Treg suppressive function in human HC Tregs and in models of colitis and GVHD (7), we propose that pretreatment with IL-27 may enhance Treg expansion and suppressive function prior to autologous Treg therapy for RRMS.

In the second scRNAseq dataset from sorted *FOXP3^+^* Tregs from RRMS patients and matched HCs, we also observed reduced expression of type I and type II IFN signaling and of Treg suppressive genes. The *in vitro* IL-27 stimulation of RRMS Tregs restored expression of multiple IFN-related genes (*IFNGR1, IRF1, IFI16*) and Treg suppressive genes (*IL7R, TIGIT, IKZF3, ICOS*).

Prior study on mice with EAE reported that Tregs within the CNS may have impaired suppressive function (59), induced by their migration through the inflamed blood-brain barrier (BBB) (17). In our *in vitro* BBB model (17), RRMS Tregs that migrated across the barrier exhibited increased *IL27R* gene expression, compared to non-migrated Tregs (Fig. 5 *E*). These results suggest that IL-27R signaling may facilitate Treg migration to the CNS, thereby enhancing their suppressive function at the sites of CNS inflammation.

## Methods

### Study Subjects

The human study was approved by the Institutional Review Board of Thomas Jefferson University. Fifty-two patients with RRMS and forty-six control subjects were enrolled after providing informed consent. Inclusion criteria for RRMS included a confirmed diagnosis of RRMS (10) and either no prior treatment with disease-modifying therapies (DMTs) or a treatment-free period of more than six months (in three patient). Control subjects had no history of inflammatory diseases and were not receiving anti-inflammatory therapy at the time of enrollment. Demographic data, disease duration, and expanded disability status scale (EDDS) scores for MS patients, as well as demographic information and diagnoses for control subjects are presented in *SI Appendix Tables S*1-3.

### Flow cytometry

Fresh peripheral blood mononuclear cells (PBMCs) from seven untreated RRMS patients and matched control subjects were stained for surface markers CD4, CD25, CD127 (for gating). The list of antibodies is provided in *SI Appendix,* Table S4.

PBMCs were isolated using Ficoll-Paque density gradient centrifugation. CD3^+^CD4^+^ cells were enriched by negative magnetic bead separation and stained for CD4, CD25, and CD127. CD3^+^CD4^+^CD25^+^CD127^low^ cells were sorted using a FACS Aria fusion cell sorter. Sorted cells were submitted for scRNAseq within 18 hours from sample collection. Mouse flow cytometry methods are described in the extended methods section in *SI Figures.* Antibodies are listed in *SI Appendix,* Table S4.

**Single-cell RNA sequencing** was performed on sorted CD4^+^CD25^+^CD127^low^ Treg cells from seven recently diagnosed, untreated RRMS patients and seven age-, sex-, and race-matched control subjects. Tregs from two MS patients and three controls were stimulated *in vitro* with IL-27 (25 ng/ml) for 1 hour, washed, and submitted for scRNseq at the Center for Applied Genomics, Children’s Hospital of Philadelphia.

### Quantitative Real-Time PCR (qRT-PCR)

The gene expression of Treg cells from 10 RMMS patients and matched controls was measured by qRT-PCR using Taqman Gene Expression Assays (Applied Biosystems) (19). Mouse RT-PCR methods are described in extended methods section in *SI Figures*. The list of human and mouse primers used in the study is provided in *SI Appendix,* Table S4.

### ELISA

CSF and serum samples from 35 RRMS patients and 29 control subjects were used to measure IFN-β and IL-27 by ELISA, as per modified manufacturer protocol (3). The sample incubation was prolonged to 24 h at 4°C, and the detection Ab incubation time was 2 h at room temperature (3). Human and mouse ELISA kits are listed in *SI Appendix,* Table S4.

### EAE

Animal experiments were conducted in accordance with protocols approved by the Institutional Animal Care and Use Committee (IACUC) at Thomas Jefferson University. Active RREAE was induced in female SJL mice (8-10 weeks old) via immunization with PLP139-151 peptide, and in BL/6 WT and Treg^ΔIL-27RA^ mice with MOG35-55 peptide. Mice with EAE received I.M. injections of cGAMP-MPs or blank-MPs (5 µg/mouse) starting either at disease onset (day 9 p.i.) or at the peak of disease (day 15 p.i.), for a total of 5 doses administered every other day.

### Migration assay

Publicly available bulk RNA-sequencing data were obtained from the Gene Expression Omnibus (GEO), accession number GSE255171. This dataset includes Treg cell transcriptomes derived from Boyden chamber migration assays of samples from 5 HCs and 4 untreated RRMS patients (uRRMS), as described by Baeten et al. (17). *IL27Ra* expression was extracted and analyzed as part of the current study. Stat analysis was performed using the two-way ANOVA test.

## Authors’ contribution

ME, MS, YW, SSG, EK, PB, and IH conducted the experiments; SMP contributed clinical data; JW, AW, MRG and BM analyzed scRNAseq data; BB, JT, BM and SMP supported the study; SMP and BM supervised the project; SMP wrote the manuscript.

## Acknowledgments

We thank the study participants, Dr. Xin Zhang (Duke University) for assistance with CSF and serum ELISA experiments, Cole J. Batty, Dr. Eric M. Bachelder, and Dr. Anslie Kristy (University of North Carolina at Chapel Hill) for providing cGAMP- and blank-MPs. We are grateful to Dr. Dhanashri Miskin (Thomas Jefferson University) for patient referral. We thank Dr. Bogoljub Ciric and Dr. Roland Martin for critical reading of the manuscript.

