## Supplementary Tables 1-4 for "Targeting Impaired Type I Interferon–IL-27 Signaling Rescues T Regulatory Cell Suppressive Function in Relapsing-Remitting Multiple Sclerosis"

**Table 1. (Study subjects for Figure 1.)**

| Subject | Age (years) | Sex | Race | Disease duration (month) | EDSS |
|---|---|---|---|---|---|
| MS1 | 39 | F | C | 2 | 2 |
| MS2 | 30 | M | AA | 3 | 1 |
| MS3 | 28 | M | C | 2 | 1 |
| **average** | **32.3** | **1F, 2 M** | **1 AA, 2 C** | **2.3** | **1.3** |

| bject | Age | Sex | Race | Diagnosis |
|---|---|---|---|---|
| HC1 | 38 | F | C | migraine |
| HC2 | 30 | M | AA | vasovagal syncope |
| HC3 | 27 | M | C | bipolar disorder |
| **average** | **31.7** | **1F, 2 M** | **1 AA, 2 C** | |

| Subject | Age (years) | Sex | Race | Disease duration (month) | EDSS |
|---|---|---|---|---|---|
| MS4 | 30 | M | AA | 3 | 1 |
| MS5 | 28 | M | C | 3 | 1 |
| MS6 | 37 | M | C | 2 | 0 |
| MS7 | 34 | F | AA | 60 | 2 |
| MS8 | 48 | M | C | 3 | 2 |
| MS9 | 52 | F | C | 1 | 0 |
| MS10 | 45 | F | AA | 120 | 0 |
| MS11 | 33 | M | C | 1 | 1 |
| MS12 | 24 | F | AA | 2 | 0 |
| MS13 | 27 | F | C | 3 | 1 |
| **average (n=10)** | **37.5** | **5 F, 5 M** | **4 AA, 6 C** | **24** | **0.8** |

| Subject | Age | Sex | Race | Control Diagnosis |
|---|---|---|---|---|
| HC4 | 30 | M | AA | vasovagal syncope |
| HC5 | 27 | M | C | bipolar disorder |
| HC6 | 38 | M | C | functional neurological disorder |
| HC7 | 34 | F | AA | migraine |
| HC8 | 48 | M | C | chronic migraine, chronic pain syndrome |
| HC9 | 52 | F | C | paresthesia, polyneuropathy |
| HC10 | 44 | F | AA | chronic migraine, chronic pain syndrome |
| HC11 | 34 | M | C | healthy donor |
| HC12 | 24 | F | AA | headache |
| HC13 | 28 | F | C | neck pain |
| **average (n=10)** | **36.7** | **5 F, 5 M** | **4AA, 6C** | |

**Table 2. (Study subjects for Figure 2.)**

| | Age (years) | Sex | Race | Disease duration (month) | EDSS | sample |
|---|---|---|---|---|---|---|
| MS14 | 43 | M | AA | 36 | 6.5 | CSF |
| MS15 | 37 | F | C | 48 | 1.5 | CSF |
| MS16 | 36 | F | C | 12 | 1 | CSF |
| MS17 | 40 | F | AA | 24 | 2 | CSF |
| MS18 | 52 | F | C | 6 | 2.5 | CSF |
| MS19 | 41 | M | AA | 12 | 3.5 | CSF |
| MS20 | 33 | F | C | 6 | 1.5 | CSF |
| MS21 | 34 | M | AA | 12 | 1 | CSF |
| MS22 | 43 | F | AA | 1 | 3 | CSF |
| MS23 | 30 | M | C | 6 | 0 | CSF |
| MS24 | 45 | F | C | 12 | 1 | CSF |
| MS25 | 27 | M | C | 0.2 | 1 | CSF |
| MS26 | 42 | F | AA | 12 | 1 | CSF |
| MS27 | 38 | F | C | 2 | 2 | CSF |
| MS28 | 50 | M | C | 12 | 2 | CSF |
| **average** | **39.4** | **9 F, 6 M** | **6 AA, 9 C** | **13.4** | **2.0** | |
| MS29 | 48 | M | AA | 6 | 5 | serum |
| MS30 | 34 | M | AA | 12 | 1 | serum |
| MS31 | 41 | F | NA | 1 | 0 | serum |
| MS32 | 43 | M | C | 18 | 2 | serum |
| MS33 | 29 | F | C | 1 | 1 | serum |
| MS34 | 30 | F | C | 2 | 1 | serum |
| MS35 | 37 | M | C | 0.6 | 2 | serum |
| MS36 | 33 | F | C | 2 | 3 | serum |
| MS37 | 33 | F | C | 12 | 2 | serum |
| MS38 | 37 | F | C | 0.6 | 1 | serum |
| MS39 | 62 | F | C | 12 | 2 | serum |
| MS40 | 47 | M | C | 6 | 2 | serum |
| MS41 | 60 | M | AA | 48 | 1 | serum |
| MS42 | 57 | F | AA | 120 | 2 | serum |
| MS43 | 34 | F | C | 2 | 3.5 | serum |
| MS44 | 49 | F | C | 5 | 2.5 | serum |
| MS45 | 39 | F | C | 0.9 | 1 | serum |
| MS46 | 38 | F | C | 1 | 0 | serum |

| | | | | | | |
|---|---|---|---|---|---|---|
| MS47 | 36 | M | C | 9 | 0 | serum |
| MS48 | 38 | F | C | 2 | 2 | serum |
| **average** | **41.3** | **13 F, 7 M** | **4 AA, 15 C, 1 NA** | **13.1** | **1.7** | |

| Subject | Age | Sex | Race | Control Diagnosis | sample |
|---|---|---|---|---|---|
| HC14 | 54 | F | C | small vessel disease | CSF |
| HC15 | 56 | F | AA | small vessel disease | CSF |
| HC16 | 40 | F | C | migraine | CSF |
| HC17 | 47 | F | C | migraine | CSF |
| HC18 | 71 | F | C | altered mental status | CSF |
| HC19 | 28 | F | C | intractable migraine | CSF |
| HC20 | 34 | F | C | chronic pain syndrome | CSF |
| HC21 | 43 | F | O | intractable migraines | CSF |
| HC22 | 41 | F | C | Bell’s palsy | CSF |
| HC23 | 65 | M | C | end stage renal disease | CSF |
| HC24 | 56 | F | AA | peripheral neuropathy | CSF |
| HC25 | 45 | F | C | diffuse myalgias | CSF |
| HC26 | 42 | F | C | chronic pain syndrome | CSF |
| HC27 | 33 | F | C | chronic pain syndrome/ restless leg syndrome | CSF |
| HC28 | 55 | F | C | peripheral neuropathy | CSF |
| HC29 | 38 | F | C | hemisensory deficit, tingling | CSF |
| HC30 | 46 | F | C | fatigue | CSF |
| HC31 | 53 | F | C | stroke | CSF |
| HC32 | 62 | F | AA | sensory polyneuropathy | CSF |
| HC33 | 29 | F | C | migraine | CSF |
| **average (n=20)** | **46.9** | **19 F, 1 M** | **3 AA, 16 C, 1 O** | | |

| | | | | | |
|---|---|---|---|---|---|
| HC34 | 46 | F | C | sleep apnea, fatigue | serum |
| HC35 | 55 | F | C | peripheral neuropathy | serum |
| HC36 | 34 | F | C | right sided numbness, carpal tunnel syndrome | serum |
| HC37 | 38 | F | C | facial and extremities tingling | serum |
| HC38 | 53 | F | C | stroke | serum |
| HC39 | 30 | F | C | urinary retention | serum |
| HC40 | 38 | F | C | fibromyalgia rheumatica | serum |
| HC41 | 34 | F | AA | headache | serum |
| HC42 | 43 | F | C | migraine | serum |
| **average (n=9)** | **41.2** | **19 F, 1 M** | **3 AA, 16 C, 1 O** | | |

**Table 3. (Study subjects for Figure 5.)**

| Subject | Age (years) | Sex | Race | Disease duration (month) | EDSS |
|---|---|---|---|---|---|
| MS49 | 49 | F | C | 6 | 3 |
| MS50 | 42 | M | C | 2 | 2 |
| MS51 | 43 | F | AA | 1 | 2.5 |
| MS52 | 36 | F | C | 84 | 1 |
| **average (n=4)** | **41.9** | 3 F, 1 M | 1 AA, 3 C | **23.3** | **2.1** |

| Subject | Age | Sex | Race | Control Diagnosis |
|---|---|---|---|---|
| HC43 | 51 | F | C | migraine |
| HC44 | 40 | M | C | subdural hematoma |
| HC45 | 44 | F | AA | migraine |
| HC46 | 38 | F | C | essential tremor |
| **average (n=4)** | **40.2** | 3 F, 1 M | 1 AA, 3 C | |

**Table 1-3.** Demographic data for MS patient and HC donors. EDSS, Expanded Disability Status Scale.

**Table 4. (Reagents used in the study)**

| Human Antibodies | | | | |
|---|---|---|---|---|
| **Antigen** | **Fluorochrome** | **Clone** | **Supplier** | **Identifier** |
| CD3 | PE-Cyanine5.5 | SK7 | Invetrogen | 35-0036-42 |
| CD4 | FITC | VIT4 | Milteny Biotec | 130-113-213 |
| CD4 | BV786 | SK3 | BD Biosciences | 563877 |
| CD25 | APC | BC96 | BioLegend | 302604 |
| CD127 | PE/Cy7 | A019D5 | BioLegend | 351320 |
| CD25 | APC | REA570 \| 3G10 | Milteny Biotec | 130-133-749 |
| CD127 | PE | REA614 \| A019D5 | Milteny Biotec | 130-113-414 |
| **Mouse Antibodies** | | | | |
| CD45 | AF700 | 30-F11 | eBioscience | 6-0451-82 |
| CD3 | BV650 | 17A2 | BioLegend | 100229 |
| CD4 | FITC | RM4-5 | eBioscience | 11-0042-82 |
| ICOS | Pe-Cy7 | 7E.17G9 | ioLegend | 117422 |
| PD-1 | PE | PC61 | BioLegend | 102008 |
| Ly6C | APC | HK1.4 | BioLegend | 128016 |
| IL-27 | APC | MM27.7B1 | BD Biosciences | 562792 |
| CD11b | PE | M1/70 | BioLegend | 101208 |
| CD11c | BV421 | N418 | BioLegend | 117329 |
| CD80 | PE | 16-10A1 | BioLegend | 104708 |
| CD86 | BV786 | GL-1 | BioLegend | 105043 |
| IL-27RA | PE | W16125D | BioLegend | 159004 |
| TGF-β | BV421 | TW7-16B4 | BioLegend | 141408 |
| Granzyme B | PE | 3G8.5 | BioLegend | 149704 |
| TNF-α | BV605 | MP6-XT22 | BD Biosciences | 569296 |
| IFN-γ | BV605 | XMG1.2 | BioLegend | 505840 |
| IL-17A | Pe-Cy7 | TC11-18H10.1 | BioLegend | 506922 |
| LAG-3 | BV421 | C9B7W | BioLegend | 125221 |
| TIGIT | BV605 | 1G9 | BioLegend | 142121 |
| GITR | BV421 | DTA-1 | BioLegend | 126331 |
| IL-10 | PE | JES5-16E3 | BD Biosciences | 554467 |
| CTLA-4 | Pe-Cy7 | UC10-4B9 | BioLegend | 106314 |
| PD-1 | BV605 | 29F.1A12 | BioLegend | 135220 |
| Ki67 | BV605 | 16A8 | BioLegend | 652413 |
| FOXP3 | APC | FJK-16s | eBioscience | 17-5773-82 |

| | | | | |
|---|---|---|---|---|
| Helios | Pe-Cy7 | 22F6 | BioLegend | 137236 |
| TIM-3 | Pe-Cy7 | RMT3-23 | BioLegend | 119716 |

| **Human Primers** | | |
|---|---|---|
| **Gene** | **Assay ID** | **supplier** |
| GAPDH | Hs02786624_g1 | ThermoFisher Scientific |
| MX1 | Hs00895608_m1 | ThermoFisher Scientific |
| ITGB7 | Hs01565750_m1 | ThermoFisher Scientific |
| ISG15 | Hs01921425_s1 | ThermoFisher Scientific |
| IFITM1 | Hs00705137_s1 | ThermoFisher Scientific |
| CD81 | Hs01002167_m1 | ThermoFisher Scientific |
| CD226 | Hs00170832_m1 | ThermoFisher Scientific |
| CD52 | Hs00174349_m1 | ThermoFisher Scientific |
| **Mouse Primers** | | |
| **Gene** | **Assay ID** | **supplier** |
| Actin B | Mm02619580_g1 | ThermoFisher Scientific |
| ISG15 | Mm01705338_s1 | ThermoFisher Scientific |
| IFITM | Mm00850040_g1 | ThermoFisher Scientific |
| OAS1A | Mm00836412_m1 | ThermoFisher Scientific |

| **Human ELISA kits** | | |
|---|---|---|
| **Kit** | **Supplier** | **Identifier** |
| Human IFNb ELISA Kit | Antigenix America | RHF842CK |
| Human IL-27 ELISA Kit | R&D systems | NBP3-06784 |
| **Mouse ELISA kits** | | |
| **Kit** | **Supplier** | **Identifier** |
| Mouse IL-10 Quantikine ELISA Kit | R&D systems | M1000B |
| Mouse IL-27 p28/IL-30 ELISA Kit - Quantikine | R&D systems | M2728 |

| **EAE Reagents** | | |
|---|---|---|
| **Kit** | **Supplier** | **Identifier** |
| PLP (139-151) | ANASPEC | AS-63912 |
| MOG (35-55) | Bio-Synthesis | CRB1000379 |
| Pertussis Toxin from B. pertussis, Lyophilized in Buffer | List Labs | 180 |
| mject™ Freund's Incomplete Adjuvant (FIA) | ThermoFisher Scientific | 77145 |
