## Supplementary Figures for "Targeting Impaired Type I Interferon–IL-27 Signaling Rescues T Regulatory Cell Suppressive Function in Relapsing-Remitting Multiple Sclerosis"

### SUPPLEMENTAL MATERIALS

### EXTENDED MATERIALS AND METHODS

#### Study Subjects

The human study was approved by the Institutional Review Board (IRB) of Thomas Jefferson University. A total of 52 RRMS patients and 46 HCs provided written informed consent prior to sample collection. Inclusion criteria for RRMS patients were a confirmed diagnosis (10) and no prior treatment with DMTs or a treatment-free period of more than six months (in three patient). Among HCs, 17 donors were age-, sex-, and race-matched to RRMS blood sample donors (Fig.*1* and *5*). CSF (n=15) and serum (n=20) samples from untreated RRMS patients, as well as CSF (n=20) and serum (n=9) de-identified samples from control donors obtained in a previous study, were used for analysis (Fig. *2)*. *SI Appendix* Table S1 presents demographic and clinical data for donors used in Fig 1; Table S2 for those in Fig 2; and Table S3 for donors whose results are presented in Fig. 5.

#### Flow Cytometry

Fresh PBMCs from three untreated RRMS patients and matched HCs were stained for surface markers CD4, CD25, CD127 (Fig. 1). Antibody information is provided in *SI Appendix* Table S4. Same marker but different attached fluorochrome staining was performed for four RRMS and matched HC donors included in Fig. 5 (26).

For mouse flow cytometry, single cell suspensions were prepared from CNS infiltrates using a 40% Percoll centrifugation gradient and from spleen tissue homogenates filtered through 70 μm cell strainer. Cells were stained for myeloid cell markers: Ly6C, CD11c, CD80 and CD86, and for Treg markers: CD3, CD4, IL27R, CTLA-4, GITR, PD1, TIGIT. Intracellular staining was performed after stimulation with PMA (50 ng/ml) and ionomycin (500 ng/ml) (Sigma-Aldrich) for 2 h, and brefeldin A (1:1000 dilution) (eBioscience) was added for an additional 3 h. Cells were then fixed, permeabilized, and stained with fluorescein-conjugated antibodies against FoxP3, LAG-3, granzyme B, perforin, IL-17A, IL-10, and TGF-β, as previously reported (60). Data acquisition was performed using a BD FACSAria Fusion flow cytometer.

Marker expression was determined in this triple-gated population with FloJo 10.6 software. The results were expressed as the percentage of gated cells positive for each marker.

**Single Cell RNA Sequencing**

scRNAseq was performed on sorted $CD4^{+}CD25^{+}CD127^{low}$ Treg cells from two independent experiments: the first with three untreated RRMS patients and three matched HCs, the second with four additional RRMS patients and matched HCs. In the second experiment, Tregs from two RRMS patients and three HCs were stimulated *in vitro* with IL-27 (25 ng/ml) for 1 h, washed, and submitted for scRNseq at the Center for Applied Genomics, Children's Hospital of Philadelphia.

Next generation sequencing libraries were prepared using the Chromium Next GEM Single Cell 3' Reagent Kits v3.1. Library quality was assessed using Agilent TapeStation for sizing (bp) and KAPA qPCR for concentration. Sequencing was performed on an Illumina NovaSeq 6000 with either an S1 100 cycle flow cell v1 or SP

100 cycle flow cell v1.5, targeting a mean depth of 20,000 reads per cell. Sequencing reads were demultiplexed and aligned to the GRCh38 reference transcriptome using the Cell Ranger pipeline (v5.0.0, 10x Genomics) to generate feature barcode matrices. Secondary analysis was conducted using the Seurat package v.4.0.2. Cells were filtered based on quality thresholds for features (200 < n < 5000) and mitochondrial content (<25%). The filtered data was normalized and scaled, and then a principal component analysis (PCA) was performed on the dataset using the most variable features to produce the clusters and subsequent UMAP projection. Violin plots depict gene expression levels. Harmony R package was used to correct the batch effects due to sequencing on different days.

*Differential gene expression analysis:* DEG were identified using FindMarkers and the FindAllMarkers functions from the Seurat R package, using the default Wilcoxon testing. Genes with Bonferroni-adjusted p value <0.05 and $\log_2$FC>0.1 (Fig. 1) or log2FC>0.25 (Fig. 5) were considered significant. For DEG across the full dataset, genes had to be expressed in at least 1% of cells. DEGs presented at Fig 1. were identified in the *FOXP3*$^+$ activated Tregs (2), while in Fig 5. in *FOXP3*$^+$ sorted Tregs.

*Gene Ontology (GO) and Pathway Enrichment Analysis:* Functional annotation of DEGs was performed using the Database for Annotation, Visualization, and Integrated Discovery (DAVID; david.abcc.ncifcrf.gov). GO term enrichment included Biological Process (BP), Cellular Component (CC), and Molecular Function (MF). Enrichment factor and a p value <0.05 were used as significance criteria. Top 10 GO terms were selected. Pathway and Gene ontology (GO) data were analyzed by Fisher's exact test followed by Benjamini-Hochberg correction as implemented in DAVID software.

**Quantitative Real-Time PCR (qRT-PCR)**

For human samples, $CD3^{+}CD4+CD127^{low}$ Tregs were sorted from PBMCs as described above.

For mouse experiments, Tregs were isolated from splenocytes of cGAMP-MP and blank-MP-treated mice using the EasySep™ Mouse $CD4^{+}CD25^{+}$ Regulatory T Cell Isolation Kit II (Stem Cell technologies), followed by RNA extraction (Zymo Research) and cDNA generation (Invitrogen).

Gene expression was measured using Taqman probes (*SI Appendix Table* 4) and run in triplicate using Quantstudio 3. Gene expression was normalized to GAPDH (mouse) or to β-actin (human). Results are presented as mean ± SEM, and statistical analysis was performed using the Mann-Whitney nonparametric test. (*$p<0.05$; **, $p<0.01$).

**ELISA:**

CSF and serum samples from RRMS patients and HCs were used for IFN-β and IL-27 measurement using ELISA as per manufacturer modified protocol (3) (*SI Appendix Table* 4).

For mice sera and culture supernatants, IL-10 and Il-27 mouse ELISA kits were used (*SI Appendix, Table* 4).

**EAE Induction and Treatment**

All animal experiments were approved by the Institutional Animal Care and Use Committee (IACUC) of Thomas Jefferson University. EAE was induced in 8-10 week-old female SJL mice (Jackson Laboratory) with proteolipid protein $(PLP)_{139-151}$ peptide (50 μg/mouse) or in C57BL/6 mice with myelin oligodendrocyte glycoprotein $(MOG)_{35-55}$ peptide (200 μg/mouse), (*SI Appendix Table* 4). Mice were immunized with a 1:1 emulsion of peptide and *M. tuberculosis* (8 mg/ml) in incomplete Freund's adjuvant. Pertussis toxin (200 ng) was administered i.p. on days 0 and 2. Mice with RREAE received I.M injections of cGAMP-MPs or blank-MPs (5 μg/mouse) starting at disease onset (day 9 p.i.) or peak of disease (day 15 p.i.) every other day for 5 doses. Clinical scores were recorded daily using the following scale: 1) limp tail, 2) hind limb weakness, 3) hind limb paralysis, 4) hind limb paralysis and forelimb weakness and 5) moribund mice (61).

To test a role of IL-27 in Treg function, we induced EAE in $Treg^{\Delta IL\text{-}27RA}$ mice and treated them with cGAMP-MPs or blank-MPs. $Treg^{\Delta Il27ra}$ mice were generated by crossing *Il27ra*$^{fl/fl}$ mice with *Foxp3*$^{CreYFP}$ mice (a kind gift from Dr. Min) (6).

**Migration assay:** Publicly available bulk RNA-sequencing data were obtained from the Gene Expression Omnibus (GEO), accession number GSE255171. This dataset includes Treg cell transcriptomes derived from Boyden chamber migration assays of samples from 5 HCs and 4 untreated RRMS patients (uRRMS), as described by Baeten et al. (17). The default DESeq2 options were used, including log fold change shrinkage. DEGs were considered only significant when the Benjamini-Hochberg adjusted P value

(FDR) was < 0.05. *IL27Ra* expression was extracted and analyzed as part of the current study. Stat analysis was performed using the two-way ANOVA test.

#### Statistical Analysis

Marker expression data (qRT-PCR and flow cytometry) (Fig. 1E, 2, 3, 4) were analyzed using non-parametric Mann-Whitney test, unless data were normally distributed, in which case unpaired t-tests were applied (Fig. 4).

Cytokine levels in CSF and serum were compared using the non-parametric Mann-Whitney test for two groups with independent samples (Fig. 2).

EAE clinical scores were analyzed using two-way ANOVA. Data are presented as mean ± SEM; $*p<0.05$, $**p<0.01$, $***p<0.001$. Statistical analysis was performed using GraphPad Prism 9 and R version 4.2.2.

#### Study Approval

The human study was approved by the IRBs of Thomas Jefferson University. All subjects signed an informed consent prior to sample collection. All animal experiments were conducted in accordance IACUC protocols at Thomas Jefferson University.

### SUPPLEMENTAL FIGURES:

#### Supplemental Figure 1.

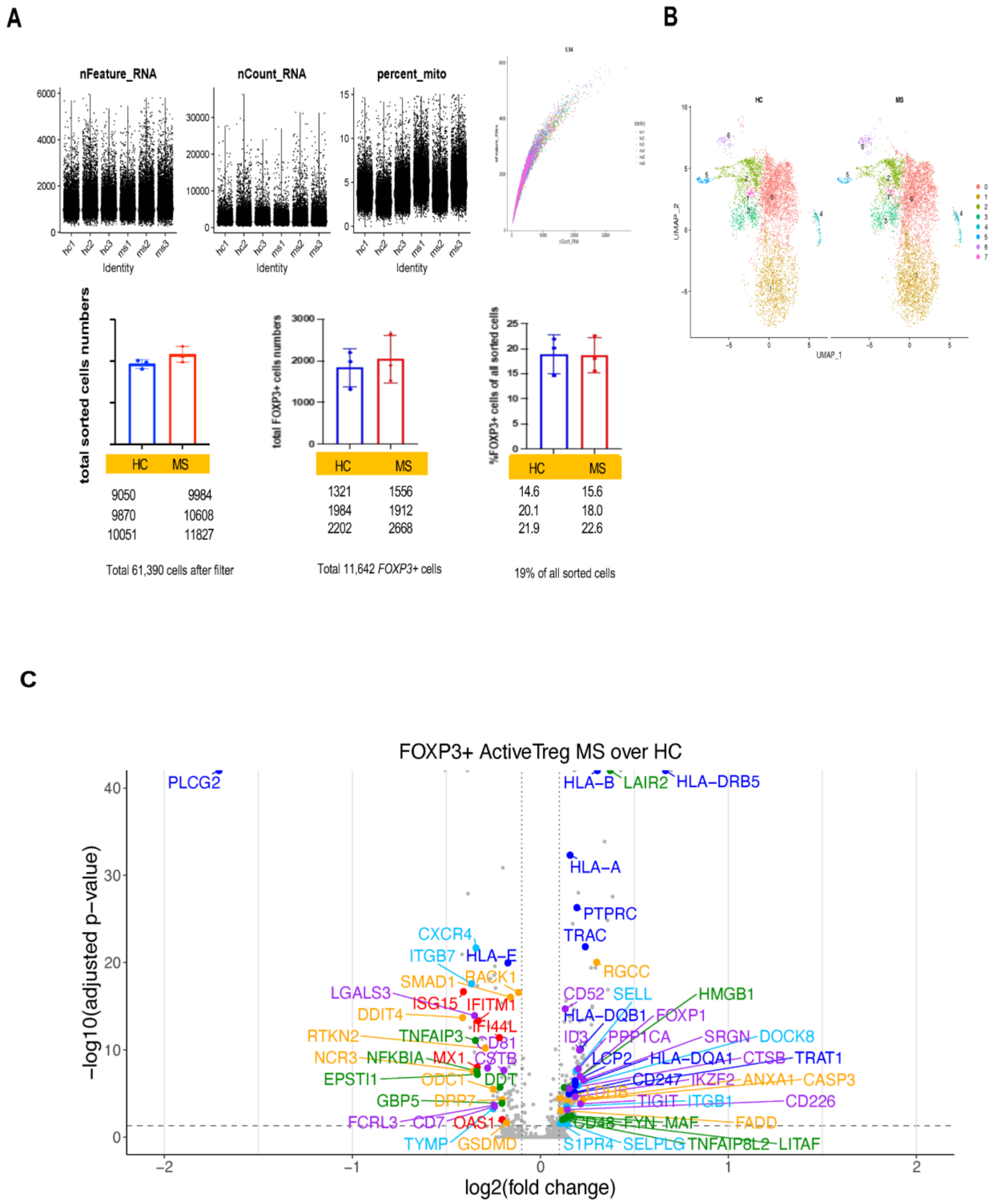

**Supplemental Figure 1.** scRNAseq quality control, filtering, and clustering of activated *FOXP3*$^+$ Treg cells. (*A*) Within Seurat, cells were filtered based on quality thresholds for features (200 < n < 5000), nCount (200<n<4000), mitochondrial content (<15%), ribosome content (>5%). A total of 61,390 cells were obtained after filter, of which 11,642 *FOXP3*$^+$ cells were analyzed. (*B*). UMAP plot of sorted Treg cells clustered in eight clusters. *(C)* Volcano plot showing biologically relevant DEGs in RRMS Tregs compared to HCs. Color codes: type I IFN (red), cell death (orange), migration (yellow), cytokine signaling (green), TCR signaling (blue) and Treg suppressive genes (purple).

**Supplemental Figure 2.**

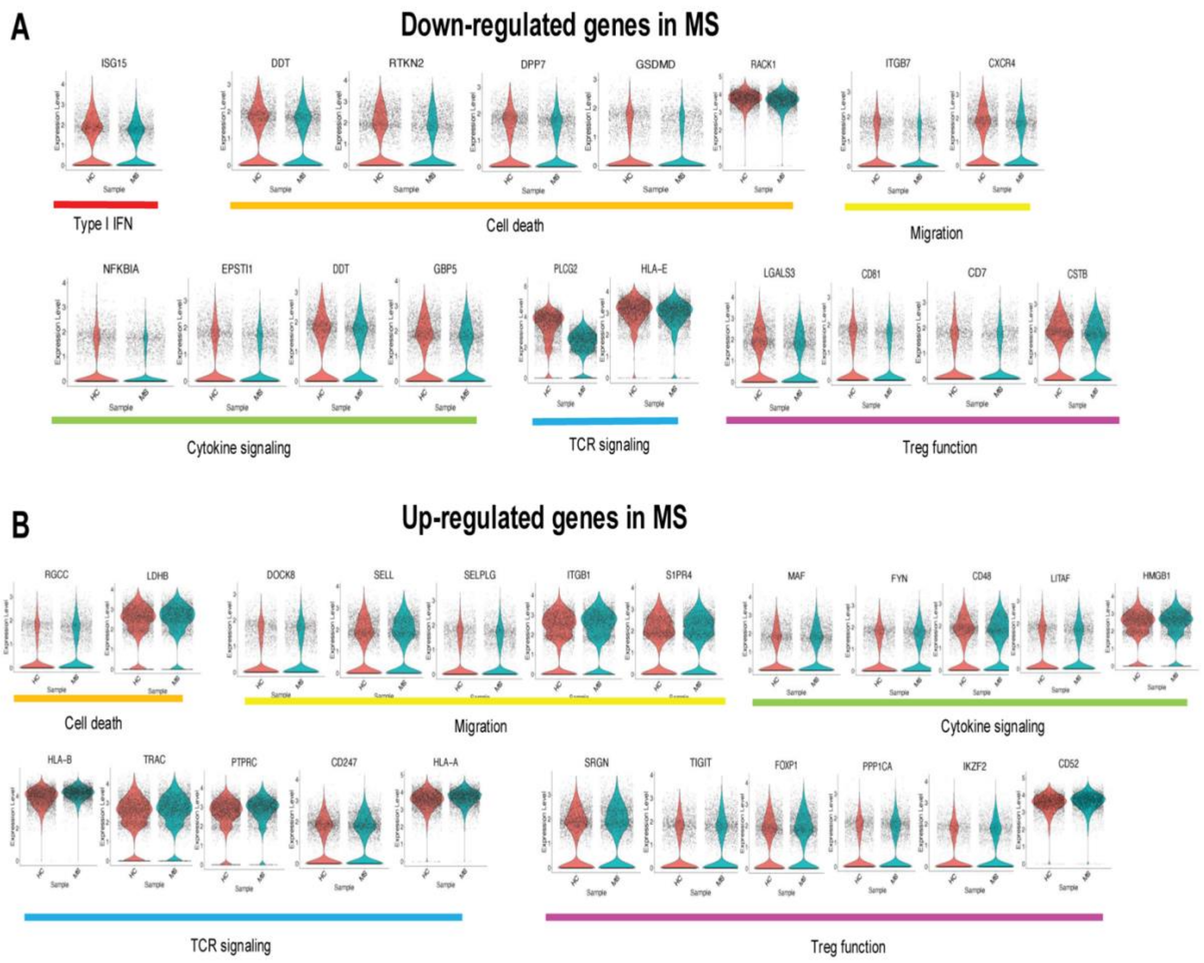


**Supplemental Figure 2.** Violin plots showing expression levels of selected biologically relevant up- and down-regulated DEGs. Manually curated DEGs (Fig. 1 *D*) are grouped in six functional categories and presented in violin plots: 1) type I IFN-related genes (red); 2) migration-related genes (yellow); 3) cell death-related genes (orange); 4) cytokine signaling (green); TCR signaling (blue), and Treg suppressive function genes (purple). *(A)* Selected down-regulated gene expression and *(B)* up-regulated genes in RRMS Tregs.

**Supplemental Figure 3.**

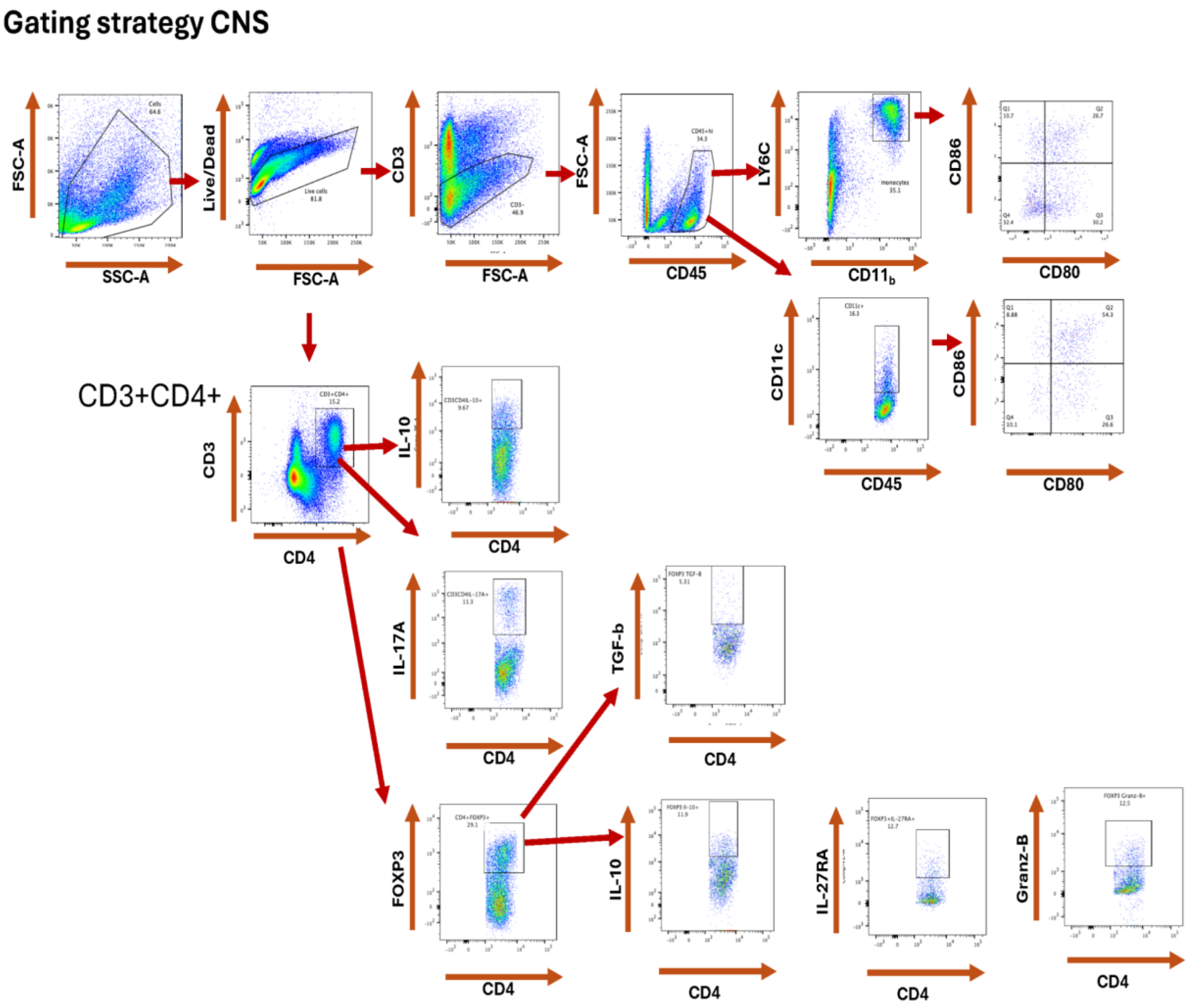


**Supplemental Figure 3.** Gating strategy for CNS-infiltrating Treg and Tconv cells. Flow cytometry gating strategy used to identify $CD3^+CD4^+FOXP3^+$ regulatory T cells (Tregs) and $CD3^+CD4^+FOXP3^-$ conventional T cells (Tconv) from CNS-infiltrating immune cells (data presented in Fig 3-4, and *SI Appendix, Fig. S*7).

**Supplemental Figure 4.**

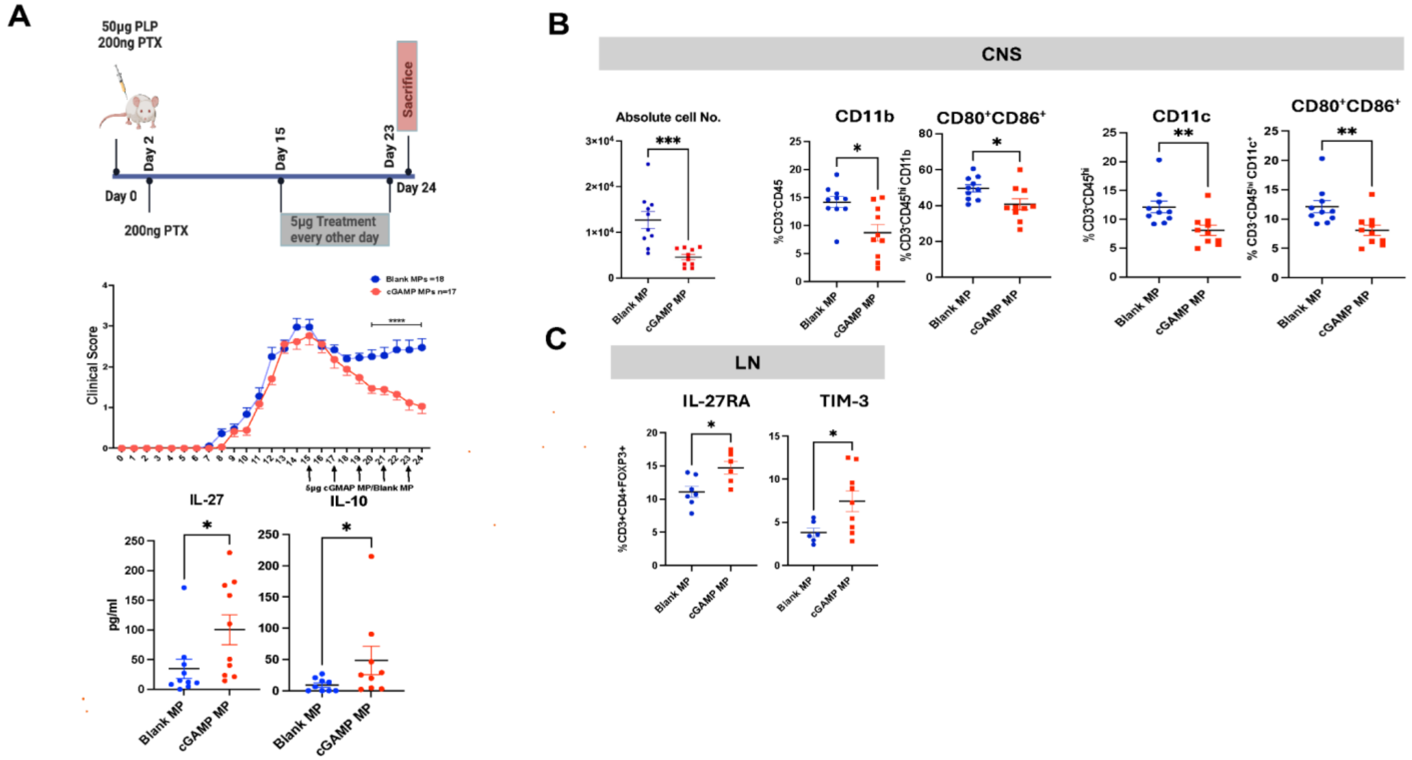


**Supplemental Figure 4.** cGAMP-MP treatment starting at the peak of disease ameliorates clinical RRAEA. (*A*). SJL/J female mice were treated with cGAMP-MPs or blank-MPs starting at the peak of disease (day 15 p.i.), 9 mice per group. Data shown are representative of two independent experiments. A significant reduction in clinical scores in cGAMP-MP-treated mice was detected by 2-way ANOVA, **** p<0.0001. Serum ELISA measurements of IL-27 and IL-10 in mice sacrificed at day 24 p.i. *(B-C)* Flow cytometry was used to evaluate the percentage of CNS-infiltrating monocytes and DCs expressing CD80 and CD86, and the expression of IL-27R and TIM-3 in gated $CD3^+CD4^+FoxP3^+$ Tregs from LN cells, Mann Whitney test (additional tested markers are shown in *SI Appendix Table S*4.

**Supplemental Figure 5**.

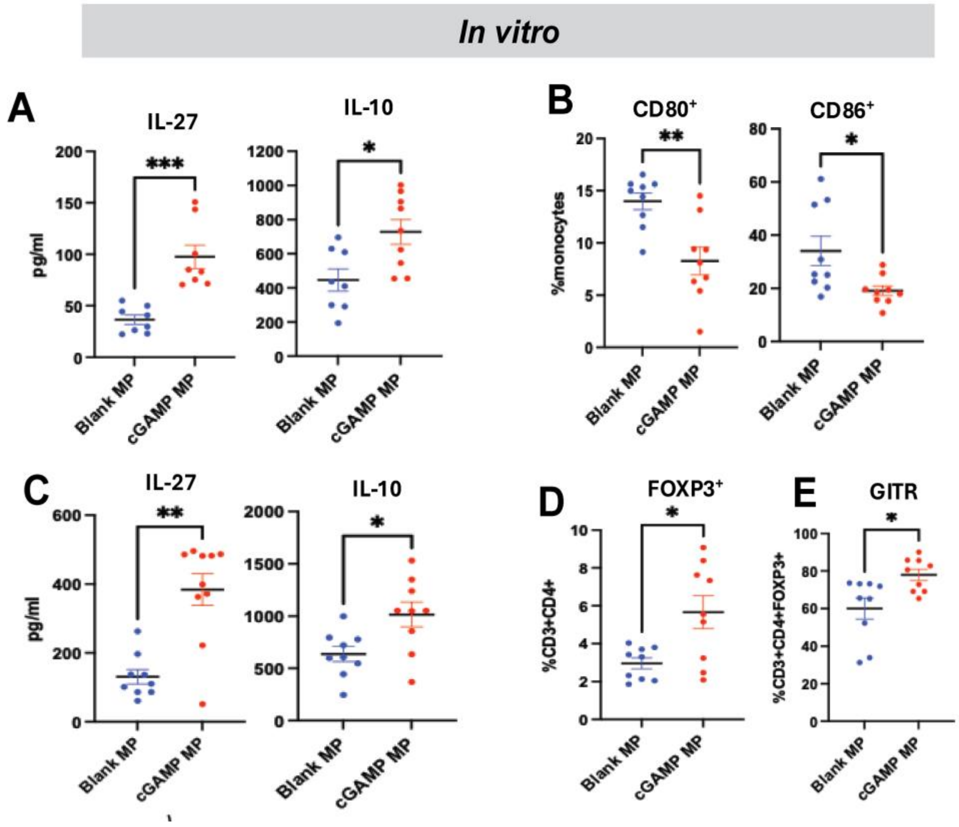


**Supplemental Figure 5**. *In vitro* co-culture of cGAMP-MP-treated monocytes with CD4$^+$ T cells induces expansion of CD3$^+$CD4$^+$Foxp3$^+$ Treg cells. *(A)* IL-27 and IL-10 was detected using ELISA in supernatants (SN) from cGAMP- and blank-MP pre-treated monocytes (48 hours)**;** *(B*) Surface expression of CD80 and CD86 was analyzed in cGAMP- and blank-MP-treated monocytes using flow cytometry**.** *(C)* IL-27 and IL-10 levels were quantified in SNs of *in vitro* cocultures of cGAMP-MP- and blank-MP-pre-treated monocytes with CD4$^+$ cells from mice with EAE (n=9 per group). *(D)* The frequency of Foxp3$^+$CD3$^+$CD4$^+$ and GITR$^+$ Tregs was assessed in CD4$^+$ cells co-cultured with pre-treated monocytes. Mann-Whitney test, * $p<0.05$, **$p<0.01$, ***$p<0.001$.

**Supplemental Figure 6.**

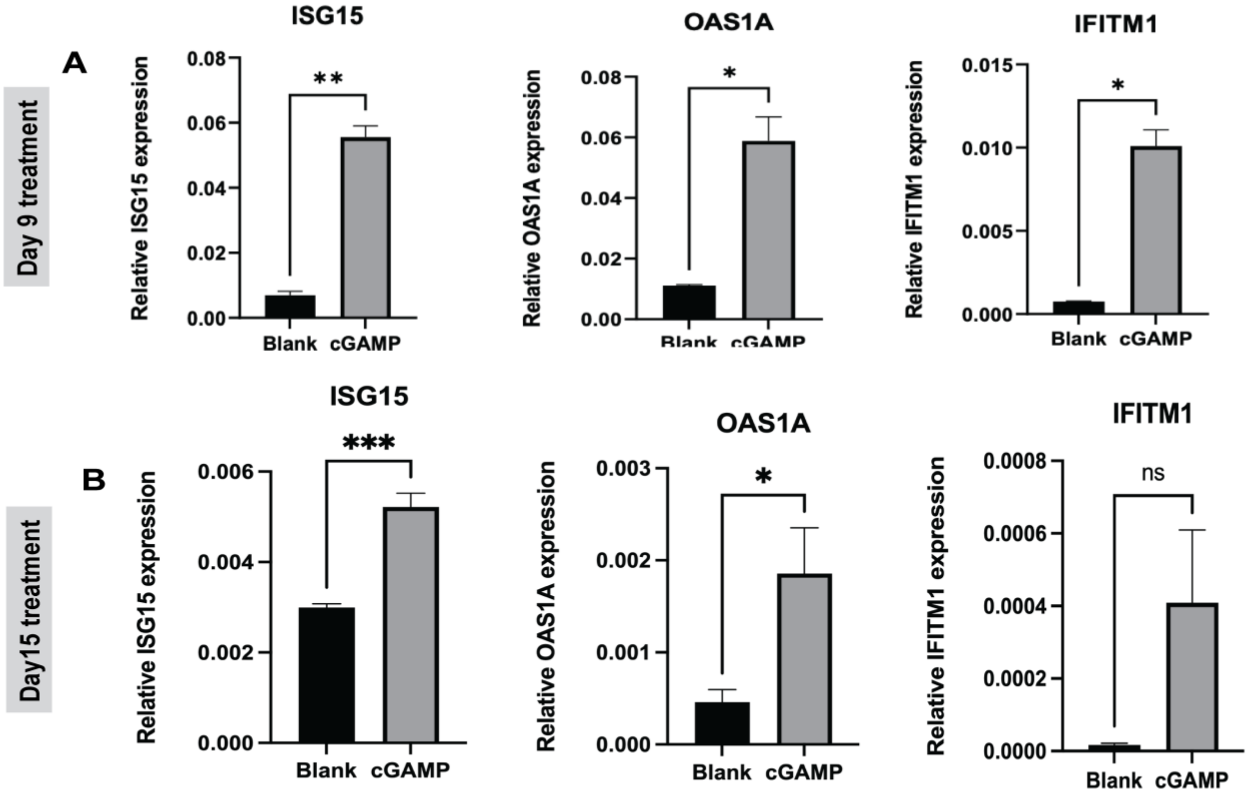


**Supplemental Figure 6.** Decreased type I IFN-related and Treg suppressive genes detected in RRMS Treg scRNAseq study were validated as increased in cGAMP MP- treated mice with EAE, suggestive of type I IFN/IL-27-induced normalization of those genes in mouse Tregs. *(A)* RT-PCR was performed on cDNA from the spleen Tregs of mice treated with cGAMP-MP or blank-MP starting at the onset of clinical EAE (day 9 p.i.) and (*B*) at the peak of the EAE clinical symptoms (day 15 p.i., 6-7 mice per group). Relative gene expression normalized against GAPDH, statistical analysis using Mann-Whitney test, * $p<0.05$, ** $p<0.01$, *** $p<0.001$.

**Supplemental Figure 7.**

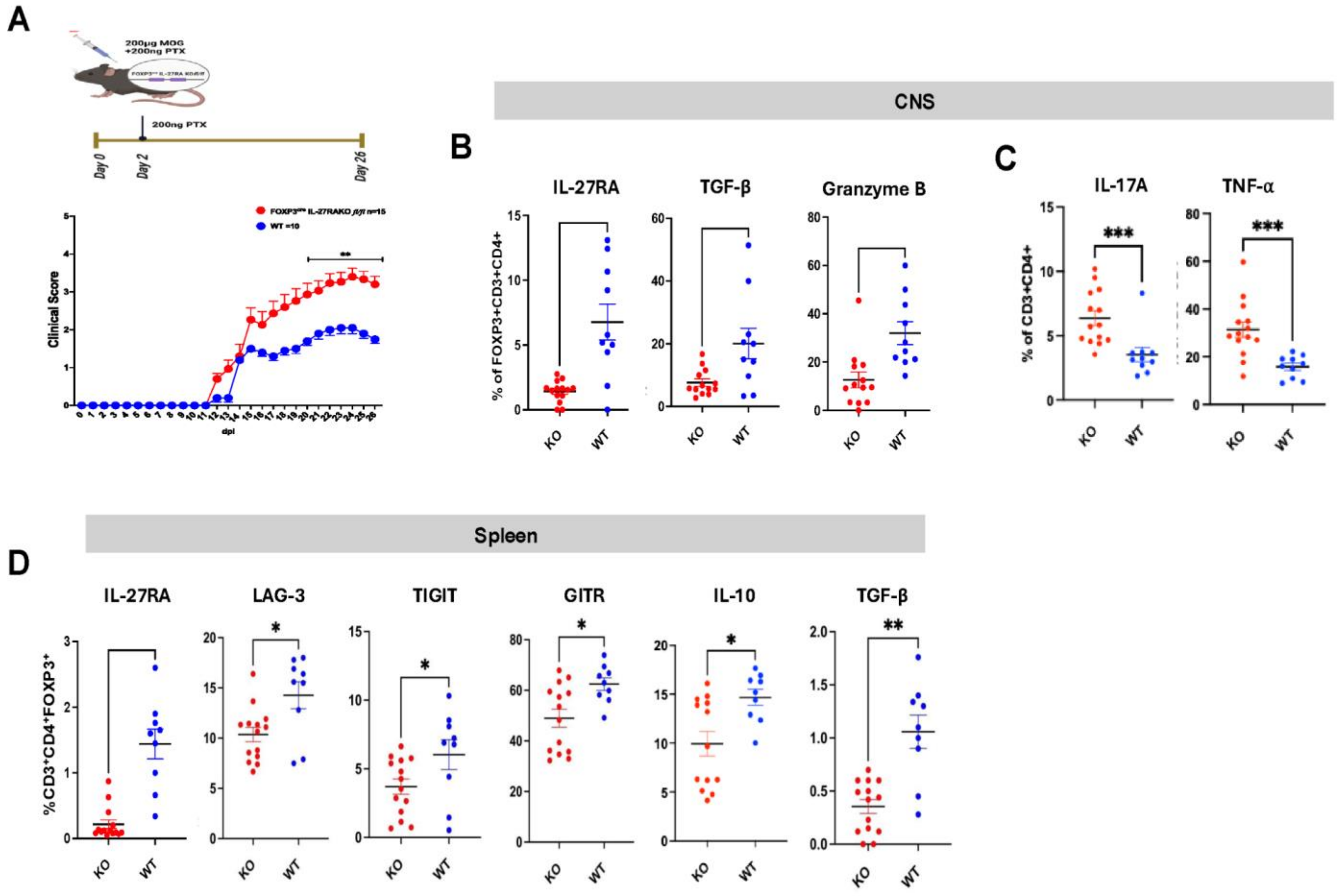


**Supplemental Figure 7.** Treg suppressive function is decreased in Treg$^{\Delta IL27ra}$ mice with EAE. *(A)* Treg$^{\Delta IL27ra}$ mice exhibited significantly higher EAE clinical scores than WT mice, indicating impaired Treg suppressive function (n=15 KO and 10 WT mice per group). Clinical scores were analyzed using two-way ANOVA. *(B, C)* Intracellular staining was used to quantify the percentage of FoxP3$^+$ Treg cells expressing anti-inflammatory cytokines IL-10, TGF-β, and Granzyme B. *(D)* Spleen Treg$^{\Delta IL27ra}$ mice-derived FOXP3$^+$Tregs were analyzed for expression of LAG-3, TIGIT, GITR, IL-10 and TGF-β. Mann-Whitney test, *p<0.05, **p < 0.01, ***p<0.001.
